# High-fidelity bispecific protease KRysin for deeper plasma proteomics and phosphoproteomics

**DOI:** 10.64898/2026.10.20.760339

**Authors:** Tanveer S. Batth, Cristina Hernandez-Rollan, Claire Koenig, Florian Liam Mao, Jesper V. Olsen

## Abstract

Bottom-up proteomics relies on efficient and reproducible proteolytic digestion. Yet trypsin, the most widely used protease, frequently misses cleavages, particularly near prolines, acidic and phosphorylated residues. Incomplete digestion inflates search space, dilutes peptide signal, and limits depth and reproducibility, especially in challenging matrices such as plasma and phosphoproteomics. Here we present KRysin, a recombinant bispecific protease that cleaves C-terminal to lysine and arginine with very high specificity and efficiency, generating tryptic peptides compatible with standard proteomics workflows. In HeLa digests, ∼99% of KRysin-generated peptides ended in lysine or arginine, including at sites followed by proline, and only 0.2% of peptides contained an internal (missed-cleaved) arginine and 6% an internal lysine, compared with 9% and 25%, respectively, for trypsin. Using narrow-window DIA at 60 samples-per-day, KRysin quantified close to 50,000 class I phosphosites, >10,000 more than trypsin-based workflows, and identified up to 40% more plasma protein groups, uniquely revealing receptors, cytokines, and growth factors. KRysin therefore provides a direct alternative to trypsin wherever digestion completeness, speed and PTM coverage are limiting.

## Introduction

Proteolytic digestion is a fundamental step in bottom-up mass spectrometry (MS)-based proteomics, where the choice of protease directly determines proteome coverage, peptide identification rates, quantitative performance and the overall success of the analysis [1]. The vast majority of proteomics workflows rely on trypsin, a pancreatic serine protease first described by Kühne in 1876 [2,3]. Trypsin cleaves C-terminal to arginine and lysine [4], and the frequency of these residues in protein sequences yields peptides of a length favorable for reversed-phase chromatography and MS analysis [5]. The resulting basic C-terminal residues also carry a positive charge that favors peptide sequencing by collisional fragmentation such as CID and HCD [6]. However, trypsin digestion requires extended incubation time and remains incomplete even after overnight digestion, leaving missed cleavages [7]. These inflate the database search space and peptide redundancy, increase sample complexity and dilute signal across several precursors for the same sequence, which lowers proteome coverage. Trypsin cleavage is further reduced when a lysine or arginine is followed by proline or flanked by aspartic or glutamic acid, and near bulky or charged post-translationally modified (PTM) residues [5,7]. Native porcine-derived trypsin remains the most prominent source of trypsin used in MS-based proteomics laboratories, but chymotrypsin inherently co-purifies with trypsin because it shares similar biochemical characteristics and must be irreversibly inactivated. Moreover, a shortcoming of native trypsin is autolysis and to prevent this, sequencing-grade porcine trypsin has to be modified by reductive methylation. Although alternative site-specific proteases such as LysC, AspN and GluC have expanded the enzymatic toolkit for bottom-up proteomics [8,9], trypsin remains the default for global proteome analysis. Its limitations underscore the need for faster and more efficient digestion that still produces tryptic peptides while improving throughput, reproducibility and PTM site coverage.

These limitations are particularly consequential in clinical samples such as blood plasma and serum, which are attractive but challenging matrices for biomarker discovery owing to a protein concentration dynamic range that spans more than ten orders of magnitude [10,11]. In large plasma cohorts, the completeness and consistency of digestion also contribute directly to the technical variance of quantitative measurements [11]. The analysis of PTMs such as protein phosphorylation on serine, threonine and tyrosine residues exemplifies a second challenge for trypsin.

Phosphorylation near lysine or arginine residues impairs tryptic cleavage [12,13]. In model peptides containing phosphorylated basophilic motifs such as the protein kinase A consensus R-X-pS/pT and R-X-X-pT, in which the arginine lies one or two residues N-terminal to the phosphosite, tryptic cleavage is strongly reduced, an effect attributed to salt-bridge formation between the phosphate group and the basic side chain [12]. Poor digestion of such sequences elevates missed-cleavage rates and splits the signal for a given phosphosite across several peptide forms, which reduces the sensitivity, accuracy and precision of phosphosite quantification and, ultimately, experimental reproducibility. Increasing the amount of trypsin only partly alleviates this problem [12].

Here we present KRysin, a recombinant high-fidelity bispecific endoprotease that cleaves C-terminal to both lysine and arginine, generating “tryptic” peptides equivalent to those produced by trypsin or trypsin/LysC (Figure 1A). KRysin combines an arginine-specific and a lysine-specific cysteine endopeptidase activity (EC 3.4.22.37 and EC 3.4.22.47) and therefore digests proteins through a catalytic mechanism distinct from that of the serine proteases used in most proteomics workflows. Arginine-specific cysteine proteases of this family have long been recognized as useful tools in protein chemistry [14], but their adoption in proteomics has been very slow. We recently described a recombinant arginine-specific enzyme of this family, Arg-C Zero, which achieves near-complete cleavage at arginine [15]. We systematically characterized KRysin proteolytic digests by liquid chromatography tandem mass spectrometry (LC-MS/MS) using the model protein bovine serum albumin (BSA), HeLa cell lysates, phosphopeptides enriched from HeLa digests, human blood plasma, and benchmarked it against trypsin and trypsin/LysC. This work extends the proteomics enzyme toolkit toward deeper coverage and higher-throughput quantitative analysis of challenging samples and PTMs.

**Figure 1.**
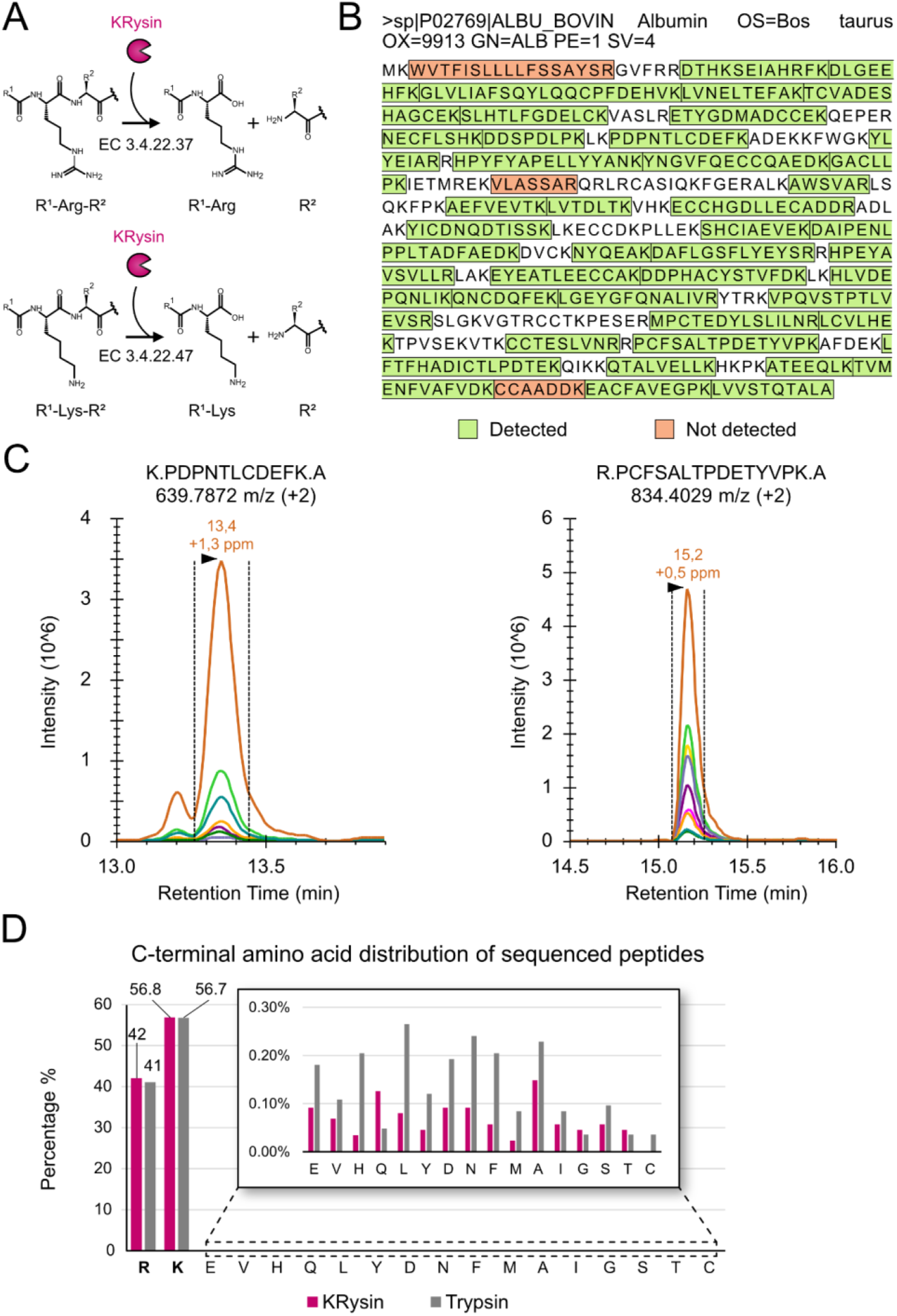
KRysin cleaves specifically after lysine and arginine, including at proline-adjacent sites, and achieves near-complete sequence coverage of BSA. (A) Reaction scheme for proteolysis catalyzed by KRysin: hydrolysis of the amide bond C-terminal to arginine (EC 3.4.22.37) and lysine (EC 3.4.22.47) residues via an active-site cysteine nucleophile, releasing a free amine on the N-terminus of the downstream fragment. (B) Sequence coverage map of bovine serum albumin (BSA, UniProt P02769) following KRysin digestion and LC-MS/MS analysis. Of 43 predicted peptides, 40 were identified (green) and 3 were not detected (orange). (C) Extracted ion chromatograms for two BSA peptides, K.PDPNTLCDEFK.A (m/z 639.7872, 2+; retention time 13.4 min, +1.3 ppm) and R.PCFSALTPDETYVPK.A (m/z 834.4029, 2+; retention time 15.2 min, +0.5 ppm), both arising from cleavage at lysine or arginine followed by proline. (D) C-terminal amino acid distribution of peptides identified by unspecific search in HeLa digests, comparing KRysin (magenta) and Trypsin (gray). Both enzymes generated peptides terminating predominantly in arginine (42% KRysin, 41% Trypsin) and lysine (56.8% KRysin, 56.7% Trypsin). Inset: expanded view of the remaining C-terminal amino acids, each present in less than 0.3% of identified peptides.

## Results

### KRysin has dual C-terminal cleavage specificity for arginine and lysine and generates “tryptic” peptides

To characterize the intrinsic cleavage specificity of KRysin independently of proteome complexity, we digested bovine serum albumin (BSA, UniProt P02769), a well-characterized single-protein substrate. Importantly, as KRysin hydrolyzes amide bonds in substrate proteins via an active-site cysteine nucleophile, a reductant like 10 mM dithiothreitol (DTT) needs to be added to the digestion mixture. Precursor (MS1) and fragment (MS2) chromatograms of “tryptic” BSA peptides were extracted from data-independent acquisition (DIA) LC-MS/MS data. Within the in-silico digestion and MS detection parameters, we detected and manually confirmed 40 of the 43 predicted fully cleaved peptides (Figure 1B, Supplementary Figure S1). One of the three undetected peptides (WVTFISLLLLFSSAYSR) spans the signal peptide, which is absent from mature albumin, and the other two (VLASSAR and CCAADDK) are short and hydrophilic. We also observed peptides in which proline occupied the P1′ position relative to lysine (K.PDPNTLCDEFK.A, residues 141 to 151) or arginine (R.PCFSALTPDETYVPK.A, residues 509 to 523), showing that KRysin cleaves lysine and arginine bonds followed by proline (Figure 1C), sites at which trypsin cleavage is inefficient [16]. To determine whether KRysin cleavage was restricted to the C-terminal side of arginine and lysine in a complex proteome, we digested a HeLa cell protein extract with KRysin or trypsin and analyzed the resulting peptides by data-dependent acquisition (DDA) LC-MS/MS. The raw MS data were then searched in MaxQuant [17] using an unspecific protease search strategy without defined protease specificity, which allowed the C-terminal residue of each sequenced peptide to be determined without bias. Arginine or lysine formed the C-terminus of 98.8% of KRysin peptides and 97.7% of trypsin peptides (Figure 1D, Supplementary Table S1). Each of the remaining C-terminal residues accounted for less than 0.3% of peptides with either enzyme, and for most residues their frequency was lower with KRysin than with trypsin, indicating that the cleavage specificity of KRysin is at least equal to that of trypsin.

### KRysin achieves near-complete digestion in complex proteomes and tolerates urea

We next benchmarked KRysin against trypsin and trypsin/LysC using HeLa cell protein extracts with DDA-based LC-MS/MS (Figure 2A). With KRysin, 93.9% of identified peptides had no missed cleavages, compared with 70.1% for Trypsin and 82.7% for Trypsin/LysC (Figure 2B). KRysin identified a majority of fully cleaved peptides, similar to Trypsin/LysC and higher than Trypsin, while producing far fewer missed-cleaved peptides (Figure 2C), so that the summed peptide signal was concentrated in fully cleaved peptides (Figure 2D). At the residue level, only 0.2% of KRysin peptides contained an internal (missed-cleaved) arginine and 6% an internal lysine, compared with 9% and 25% for trypsin and 9% and 9.5% for trypsin/LysC (Supplementary Figure S2). Adding LysC to trypsin thus improved cleavage at lysine but not at arginine, whereas KRysin increased cleavage efficiency at both residues. Because many sample preparation workflows require protein denaturation before or during digestion, we also evaluated KRysin across a urea dilution series from 0 to 3 M (Supplementary Table S2). KRysin remained active throughout this range. The number of fully cleaved peptides declined gradually from 10,904 at 0 M to 6,225 at 3 M urea (Figure 2E), whereas the proportion of fully cleaved peptides remained at 89% or higher up to 2.5 M and was 82% at 3 M (Figure 2F). By comparison, LysC from *Achromobacter lyticus* displayed a marked increase in missed cleavages above 1.5 M urea [18]. These results support the use of KRysin in urea-based in-solution digestion protocols with limited dilution.

**Figure 2.**
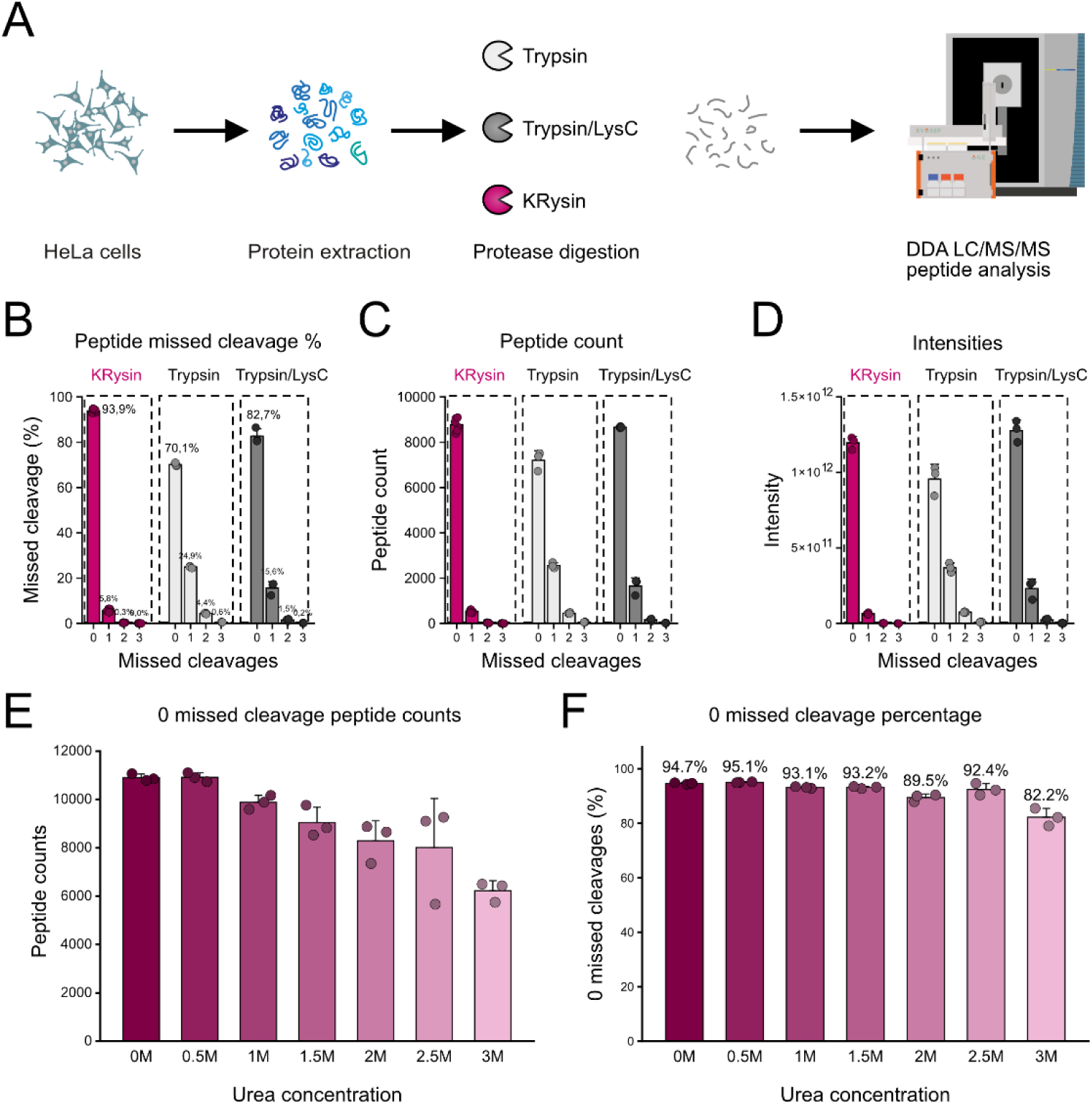
KRysin digestion of HeLa cell proteomes and tolerance to urea. (A) Workflow. Proteins extracted from HeLa cells were digested with Trypsin, Trypsin/LysC or KRysin and analyzed by DDA LC-MS/MS. (B) Percentage of identified peptides with 0 to 3 missed cleavages for each protease, with mean values labelled. (C) Unique peptide counts grouped by number of missed cleavages. (D) Summed peptide intensities grouped by number of missed cleavages. (E) Number of peptides with no missed cleavages identified after KRysin digestion in 0 to 3 M urea. (F) Percentage of peptides with no missed cleavages after KRysin digestion in 0 to 3 M urea, with mean values labelled. Bars show the mean of n = 3 digestions, error bars show SD, and dots show individual replicates.

### KRysin improves phosphosite identification by reducing missed cleavages near phosphorylation sites

Phosphorylation adjacent to lysine or arginine is a well-documented source of missed cleavages in tryptic digests [12,13], producing long, heavily missed-cleaved phosphopeptides that are difficult to detect and site localize, inflating the database search space. State-of-the-art phosphoproteomics based on trypsin-digestion workflows allow for identification of up to 40,000 localized phosphorylation sites in single-shot LC-MS analysis of a human cell line [19]. To test if KRysin can improve phosphoproteome coverage, we compared KRysin with trypsin and trypsin/LysC for the digestion of HeLa lysates at enzyme-to-protein ratios of 1:50 and 1:150, followed by automated phosphopeptide enrichment and narrow-window data-independent acquisition (nDIA) [20] on an Orbitrap Astral mass spectrometer (Figure 3A). KRysin yielded 40,537 (1:50) and 41,726 (1:150) phosphopeptides, 1-7% more than trypsin or trypsin/LysC (Figure 3B). The gain in localized class I phosphosites was considerably larger, with averages of 49,888 and 48,807 sites per sample at 1:50 and 1:150, corresponding to >10,000 more phosphosites than trypsin and trypsin/LysC, with more sites consistently identified across replicates (Figure 3C). This was accompanied by markedly improved digestion of phosphorylated peptides. At 1:50, 80.4% of phosphopeptides had no missed cleavages with KRysin, compared with 51.0% for trypsin and 57.5% for Trypsin/LysC (Figure 3D). Non-phosphorylated peptides from the same digests showed the same trend, with 93.1% fully cleaved peptides for KRysin, 86.9% for Trypsin/LysC and 80.3% for Trypsin at 1:50 (Supplementary Figure S3). Notably, KRysin at 1:150 quantified more phosphosites (48,807) than trypsin/LysC at 1:50 (39,158), with a lower missed-cleavage rate, while using one third of the enzyme amount. MS2 signal intensities were comparable across all conditions (Supplementary Figure S4), indicating that the gains did not originate from differences in MS signal.

**Figure 3.**
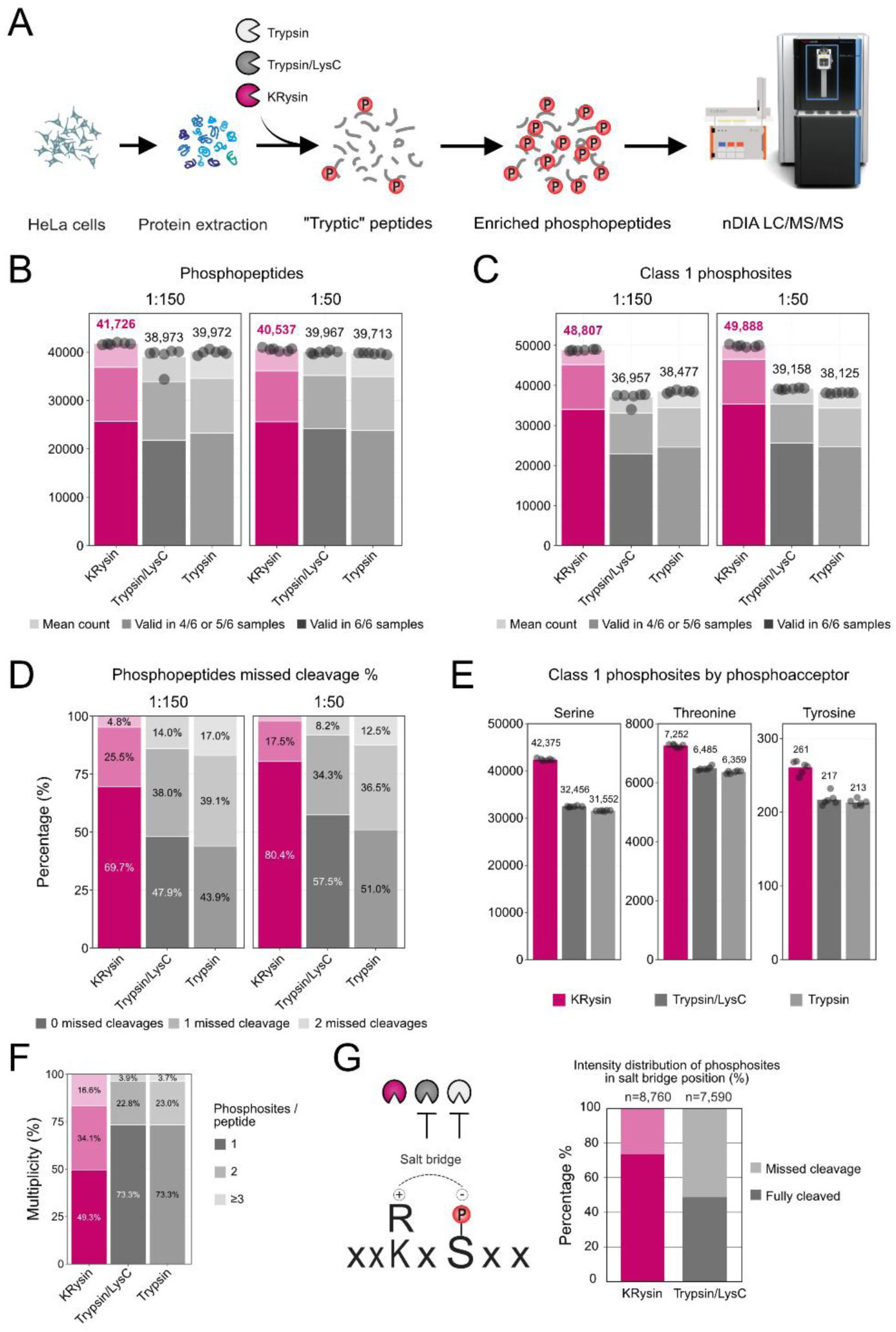
KRysin digestion improves phosphosite identification. (A) Workflow. HeLa proteins were digested with Trypsin, Trypsin/LysC or KRysin, phosphopeptides were enriched, and samples were analyzed by nDIA LC-MS/MS. (B) Phosphopeptide and (C) class I phosphosite identifications at enzyme-to-protein ratios of 1:150 and 1:50. Stacked bars show identifications valid in 6/6 samples, in 4/6 or 5/6 samples, and the mean count per sample, with mean values labelled and dots showing individual replicates. (D) Percentage of phosphopeptides with 0, 1 or 2 missed cleavages. (E) Class I phosphosites by phosphoacceptor residue (serine, threonine, tyrosine) at an enzyme-to-protein ratio of 1:50, with mean values labeled and dots showing individual replicates. (F) Phosphosite multiplicity at an enzyme-to-protein ratio of 1:50, shown as the percentage of phosphopeptides carrying 1, 2 or ≥3 phosphosites. (G) Schematic of a phosphosite in a salt-bridge position relative to an adjacent lysine or arginine, and the percentage of intensity for these phosphosites from fully cleaved versus miscleaved phosphopeptides in KRysin and Trypsin/LysC digests.

The additional phosphosites obtained with KRysin comprised serine, threonine and tyrosine sites (Figure 3E, Supplementary Figure S5A), without a change in the relative distribution of phospho-acceptor residues (Supplementary Figure S5B). Part of the larger gain in phosphosites than in phosphopeptides was explained by site multiplicity, as 50.7% of KRysin phosphopeptides identified carried two or more phosphosites, compared to only 26.7% for trypsin/LysC and trypsin (Figure 3F). Multiplicity did not account for the full gain, however, since KRysin also quantified more unique phosphosite positions than trypsin/LysC across all replicates (29,327 versus 24,405 at 1:50). Sites quantified with the two digestion strategies overlapped substantially, but a considerable fraction was unique to each, with different multiplicity profiles (Supplementary Figure S6). Sites unique to Trypsin/LysC were mostly detected on missed-cleaved phosphopeptides with short distances between cleavage sites, whereas sites unique to KRysin were found on longer, more hydrophilic peptides with larger distances between lysine and arginine residues (Supplementary Figure S7). Motif analysis showed that KRysin preferentially quantified sites in acidic regions rich in aspartic and glutamic acid, whereas Trypsin/LysC covered more proline-directed and basophilic motifs (Supplementary Figure S8). Finally, for phosphosites positioned to form a salt bridge with a lysine or arginine two residues away, 73.88% of the measured intensity came from fully cleaved phosphopeptides with KRysin, compared with 48.9% with Trypsin/LysC (Figure 3G).

### KRysin enables high-throughput plasma proteome analysis with increased depth

Blood plasma is one of the most clinically valuable sample types in proteomics, offering a minimally invasive window into systemic physiology that makes it central to biomarker discovery and diagnostics [11]. It is also a demanding biological matrix, owing to a protein concentration dynamic range of more than ten orders of magnitude and substantial inter-individual variability [10,11]. Blood plasma contains high levels of active serine proteases, such as those in the coagulation, complement and fibrinolytic cascades [21]. We next evaluated KRysin for the in-solution digestion of human donor platelet-poor plasma (Figure 4A). In an initial DDA analysis on an Orbitrap Exploris 480 mass spectrometer, 97% of the unique peptides identified from KRysin-digested plasma had no missed cleavages, compared with 82% for Trypsin/LysC and 69% for Trypsin, and KRysin yielded twice as many fully cleaved peptides as Trypsin (1,341 versus 654; Supplementary Figure S9). Accordingly, more plasma protein groups were identified with KRysin by both DDA and DIA on the same instrument (165 and 260 protein groups, compared with 134 and 209 for Trypsin/LysC and 126 and 148 for Trypsin; Figure 4B).

**Figure 4.**
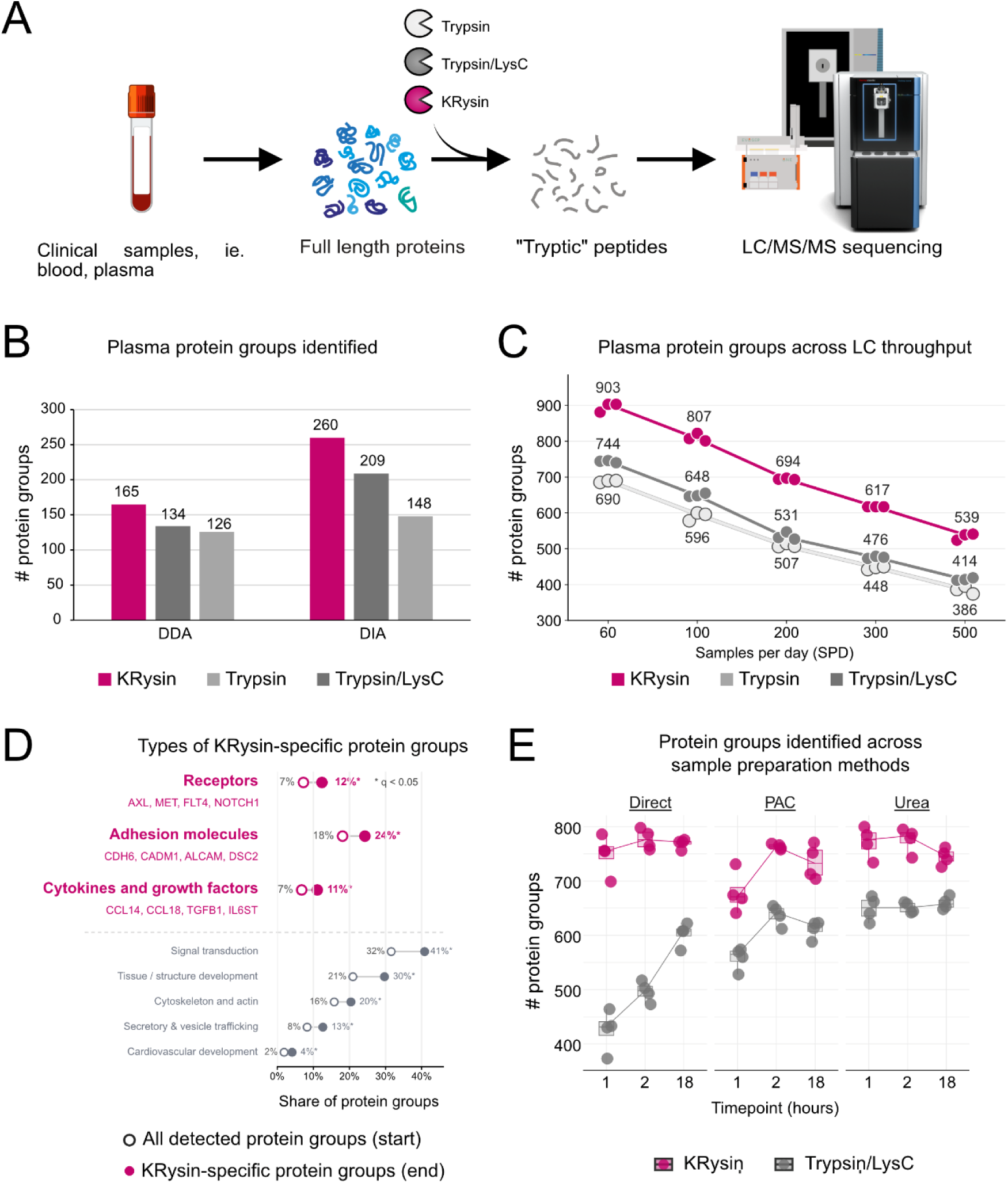
KRysin digestion increases proteome depth in human plasma. (A) Workflow. Human plasma proteins were digested with Trypsin, Trypsin/LysC or KRysin and analyzed by LC-MS/MS. (B) Plasma protein groups identified by DDA and DIA on the same instrument. (C) Plasma protein groups identified by nDIA at LC throughputs from 60 to 500 samples per day (SPD). Dots show triplicate digestions, lines connect medians, and median values are labelled. (D) Gene Ontology enrichment among the 473 KRysin-specific protein groups, shown as share of protein groups versus all detected plasma protein groups (g:Profiler, Benjamini-Hochberg FDR). Representative proteins are listed under the highlighted categories, and asterisks mark q < 0.05. (E) Protein group identifications after direct in-solution, PAC or urea-based sample preparation with 1, 2 or 18 h digestion, analyzed by nDIA on an Orbitrap Astral. Dots show individual replicates, and boxes show the median and interquartile range.

We then examined plasma proteome depth as a function of chromatographic throughput, measured as samples per day (SPD), using nDIA on an Orbitrap Astral [20]. Across throughputs from 60 to 500 SPD, KRysin identified more plasma protein groups than trypsin and trypsin/LysC at every LC gradient length, with median counts of 903 protein groups at 60 SPD and 539 at 500 SPD (Figure 4C). This corresponded to 31 to 40% more protein groups than trypsin and 21 to 31% more than trypsin/LysC, with the gain over trypsin increasing at higher throughput. As a result, KRysin at 200 SPD identified as many protein groups (694) as Trypsin at 60 SPD (690), enabling three times higher throughput using KRysin while preserving plasma proteome depth. Moreover, KRysin at 100 SPD (807) exceeded Trypsin/LysC at 60 SPD (744). To characterize the additional depth, we defined 473 KRysin-specific protein groups, identified in all three KRysin replicates but not with trypsin or trypsin/LysC at any throughput, and tested whether specific functional categories made up a larger share of this set than of all detected plasma protein groups (Gene Ontology enrichment; g:Profiler g:GOSt; Benjamini-Hochberg FDR; Figure 4D). Receptors (signaling receptor activity, GO:0038023) accounted for a larger proportion of the KRysin-specific set than of the background (12% versus 7%, q = 0.00095), including AXL, MET, FLT4 and NOTCH1. Adhesion molecules (cell junction, GO:0030054) showed the same pattern (24% versus 18%, q = 0.0033), including CDH6, CADM1, ALCAM and DSC2, as did cytokines and growth factors (cellular response to cytokine stimulus, GO:0071345; 11% versus 7%, q = 0.0099), including CCL14, CCL18, TGFB1 and IL6ST. Broader functional categories were also significantly enriched, including signal transduction (41% versus 32%), tissue and structure development (30% versus 21%), cytoskeleton and actin (20% versus 16%), secretory and vesicle trafficking (13% versus 8%) and cardiovascular development (4% versus 2%) (Figure 4D).

We further evaluated KRysin in three plasma sample preparation strategies: direct in-solution digestion in HEPES buffer without denaturation, protein aggregation capture (PAC) on magnetic beads [22], and urea-based denaturation followed by in-solution digestion, each digested for 1, 2 or 18 hours and analyzed by nDIA on an Orbitrap Astral. Across all three workflows and time points, KRysin identified more protein groups than Trypsin/LysC (Figure 4E). KRysin reached close to its maximum depth after 1 hour digest in the direct and urea workflows and after 2 hours with PAC, whereas trypsin/LysC in the direct workflow required overnight digestion and still identified fewer protein groups. These differences mirrored the missed-cleavage rates. After 1 hour, 10.7 to 14.7% of KRysin peptides contained a missed cleavage across the three workflows, compared with 26.6 to 36.7% for Trypsin/LysC (Supplementary Figure S10). Peptides identified with KRysin were predominantly fully cleaved (Supplementary Figure S11), consistent with the concentration of signal into fewer, fully cleaved peptide forms observed in the other sample types. This suggests that KRysin is ideally suited for high-throughput proteomics workflows, where shorter processing time is a key factor for scalability.

Protein quantification was reproducible with both enzymes. Without normalization, median protein CVs were lower with KRysin than with trypsin/LysC in seven of nine matched conditions, with the largest difference after direct digestion for 18 hours (9.3% versus 41.5%). In the two conditions where KRysin CVs were higher (PAC at 1 hour and urea at 2 hours), KRysin nonetheless identified more protein groups than trypsin/LysC (Supplementary Figure S12A). Much of this variation reflected differences in total signal between replicate digests, which spanned up to 1.9-fold with KRysin and up to 2.7-fold with trypsin/LysC (Supplementary Figure S12C). After median normalization, median CVs were 6 to 13% for both enzymes, with KRysin remaining lower in the direct workflow and comparable to trypsin/LysC in the PAC and urea workflows (Supplementary Figure S12B). For both enzymes, CVs were generally lowest for the most abundant proteins, and the differences between the enzymes seen in the overall CVs held across most of the abundance range (Supplementary Figure S13). After direct digestion for 18 hours, for example, the most abundant third of protein groups had a median CV of 5.5% with KRysin compared with 41.6% with trypsin/LysC. To assess whether these observations depended on the mass spectrometer or acquisition method, we analyzed the same samples on an Orbitrap Exploris 480 using DDA. Although identification numbers were lower than with nDIA on the Orbitrap Astral, KRysin again yielded more protein groups, a higher proportion of peptides without missed cleavages, and comparable or lower protein CVs than trypsin/LysC (Supplementary Figure S14).

## Discussion

The development of KRysin addresses a longstanding bottleneck in proteomics: the incomplete and inconsistent digestion of proteins by trypsin, particularly at sites adjacent to proline, acidic, or phosphorylated residues. This limitation has historically constrained the depth and reproducibility of proteomic analyses, especially in complex biological matrices such as human plasma. With recombinant KRysin, which hydrolyzes amide bonds C-terminal to lysine and arginine residues via an active-site cysteine nucleophile, this study delivers a solution that not only matches but surpasses the performance of trypsin, thereby expanding the analytical capabilities of mass spectrometry-based proteomics. Building on decades of work that have sought to overcome the shortcomings of trypsin, including the adoption of alternative proteases such as Lys-C, Lys-N and Asp-N [8,9], KRysin represents a significant extension of the proteolytic toolkit. Previous studies demonstrated that multi-protease strategies can increase proteome coverage and PTM detection [8], but each alternative enzyme brought its own limitations in specificity, efficiency, or compatibility with standard workflows. KRysin distinguishes itself by achieving near-complete cleavage at both lysine and arginine residues, including those followed by proline, a known challenge for trypsin [16]. In HeLa cell digests, KRysin generated peptides with approximately 99% ending in lysine or arginine, with missed cleavages (an internal arginine or lysine) in only 0.2% and 6% of KRysin peptides, respectively, compared to 9% and 25% for trypsin. This remarkable specificity and efficiency directly address the gaps left by previous proteases and protease combinations. These improvements translate into deeper and more reproducible measurements. In phosphoproteomics, KRysin enabled the quantification of nearly 50,000 localized class I phosphosites per sample. This is over 10,000 more than achieved with trypsin or trypsin/LysC while increasing the proportion of fully cleaved phosphopeptides from about 50% (trypsin) to 80% (KRysin). In plasma proteomics, KRysin identified up to 40% more protein groups than trypsin or trypsin/LysC, including low-abundance cell-surface receptors, adhesion molecules, cytokines, and growth factors. These advances are critical for comprehensive biomarker discovery, systems biology, and the study of signaling pathways, as they enable the detection and quantification of proteins and post-translational modifications that were previously inaccessible. By providing a more complete and reproducible proteolytic digestion process, KRysin will facilitate future proteomics research across a broad spectrum of applications. Its compatibility with high-throughput workflows supporting 60 to 500 samples-per-day LC-MS analyses makes it ideally suited for large-scale studies, clinical cohorts, and time-sensitive analyses. The ability to generate tryptic peptides that are fully compatible with existing mass spectrometry pipelines ensures that KRysin can be seamlessly integrated into current and emerging proteomics platforms, accelerating the pace of discovery.

Looking forward, the introduction of KRysin is characteristic of a broader trend in proteomics towards deeper, more comprehensive, and clinically relevant analyses. As the field moves toward routine deep proteome and PTM-resolved studies, innovations in protease engineering, mass spectrometry sensitivity, and bioinformatics integration will be essential. The capacity to achieve near-complete amino acid coverage, as demonstrated by multi-protease workflows [8], is now within reach, and KRysin sets a new standard for what is possible in both research and translational settings. Technologically, KRysin’s recombinant cysteine protease design represents a significant departure from the serine protease chemistry of trypsin. This unique enzymatic profile not only enhances digestion efficiency but also ensures robustness and scalability across diverse sample types and throughput requirements. KRysin’s adaptability and performance under mild denaturing conditions further underscore its practical utility. The integration of KRysin into high-throughput proteomics pipelines has already facilitated new research by reducing digestion times and increasing sample throughput without compromising analytical depth or accuracy. This capability is particularly valuable for studies requiring rapid turnaround and for clinical applications where sample volume and processing time are limiting factors.

Nonetheless, certain limitations remain. KRysin’s optimal performance was observed under mild denaturing conditions, with reduced activity in high urea concentrations, suggesting some constraints in compatibility with stronger denaturing protocols. Additionally, the study did not evaluate KRysin’s compatibility with workflows involving reductant-sensitive steps, which may restrict its use in specific applications. Several questions remain unanswered, including KRysin’s performance across a broader range of sample types, its efficacy in digesting proteins with diverse post-translational modifications, long-term stability, and cost-effectiveness relative to traditional proteases. Addressing these questions will be important for fully realizing the potential of this technology. Ultimately, KRysin’s ability to deliver deeper and more accurate proteomics and phosphoproteomics data from plasma and other complex samples will be invaluable for medical research. By enabling more comprehensive and reliable protein profiling, KRysin will enhance biomarker discovery, disease mechanism studies, and the development of precision medicine approaches, thereby advancing clinical practice.

## Materials and Methods

### Reagents and enzymes

KRysin (KPL ApS, Copenhagen, Denmark) was supplied lyophilized and reconstituted in milliQ water at a working concentration of 0.5 mg/mL immediately before use and stored at −20 °C. Trypsin (Sigma-Aldrich, sequencing grade) and LysC (Fujifilm Wako) were also reconstituted at a concentration of 0.5 mg/mL and were used as comparison enzymes. For Trypsin/LysC digestions, both enzymes were added simultaneously at the ratios given for each experiment. HEPES, DTT, TCEP, chloroacetamide, urea, SDS, acetonitrile, ethanol, formic acid, trifluoroacetic acid and glycolic acid were obtained from Sigma-Aldrich. SiMAG-Silanol beads (Chemicell GmbH) were used for PAC digestion experiments and Zr-IMAC HP beads (ReSyn Biosciences) were used for phosphopeptide enrichment. Enzyme-to-substrate ratios are given for each experiment below.

### Protein substrates and samples

#### Bovine serum albumin

Bovine serum albumin (BSA, Fraction V, ≥96% purity, Sigma-Aldrich #A7906) was used as a model substrate to assess KRysin cleavage specificity and sequence coverage.

#### HeLa cell culture and lysis

Human cervical carcinoma cells (HeLa) were cultured in DMEM (Gibco) containing 2 mM L-glutamine, 10% fetal bovine serum (Gibco), 100 U/mL penicillin and 100 µg/mL streptomycin at 37 °C with 5% CO_2_, manually harvested, washed with ice-cold PBS, and lysed in boiling lysis buffer (4% SDS, 100 mM Tris-HCl pH 8.5, 5 mM TCEP, 10 mM chloroacetamide) at 95 °C for 10 min with agitation, followed by probe sonication [22]. Protein concentration was determined by tryptophan fluorescence [23].

#### Human plasma

Samples were collected from surplus material from anonymous healthy adult blood donors at the Capital Region Blood Bank at the Copenhagen University Hospital-Rigshospitalet following written and informed consent. The anonymized sample collection was approved by the local ethics committee (Reference: F-24061507). Plasma was collected in 4 mL K2-EDTA tubes (Greiner Bio-One) and centrifuged for 6 min at 3000g for platelet-poor plasma (PPP). The 6 min centrifugation time corresponded to 10 min centrifugation in 2.5 cm longer 6 or 9 mL tubes. All samples were processed within 2 h after collection and frozen at −80 °C until further analysis. Separate aliquots were prepared for each workflow to avoid freeze–thaw cycles. The plasma protein concentration was approximated to 70 µg/µL.

### Protein digestion

#### BSA digestion (Figure 1B and 1C; Supplementary Figure S1)

Bovine serum albumin (BSA, 10 µg) was denatured in 8 M urea for 10 min at room temperature. The sample was then diluted to a final urea concentration of 1 M with 50 mM HEPES, pH 8.5, containing 10 mM dithiothreitol (DTT). No alkylation was performed. KRysin was added at an enzyme-to-protein ratio of 1:50 (w/w), and digestion was carried out for 2 h at 37 °C. Digestion was quenched by acidification with trifluoroacetic acid (TFA) to a final concentration of 1%, and 50 ng of peptides was loaded onto Evotips for LC-MS/MS analysis.

#### HeLa digestion for proteome benchmarking and unspecific search (Figures 1D and 2A to 2D; Supplementary Figure S2; Supplementary Table S1)

HeLa protein extracts were digested with KRysin, Trypsin or Trypsin/LysC using protein aggregation capture (PAC) on magnetic microparticles as previously described [22]. For each digestion, 10 µg of protein were aggregated on 2 µl of SiMAG-Silanol magnetic beads by the addition of acetonitrile (ACN) to a final concentration of 70%. The beads were washed once with 100% ACN and once with 70% ethanol, followed by on-bead digestion in buffer containing 50 mM HEPES pH 8.5 with the addition of 10 mM DTT for the digestions with KRysin. Digestion was performed using a 1:50 enzyme-to-protein ratio and the digestion was carried out overnight at 37 °C with shaking, subsequently quenched by the addition of 1% final concentration of TFA. Peptides corresponding to 500 ng were loaded onto Evotips for LC-MS/MS analysis. Digestions were performed in triplicate.

#### Urea tolerance (Figure 2E and 2F; Supplementary Table S2)

HeLa lysate (10 µg, in SDS-containing lysis buffer) was processed by PAC as described above, which also removed the SDS. After the washes, beads were resuspended in 50 mM HEPES, pH 8.5, containing 10 mM DTT and urea at 0, 0.5, 1, 1.5, 2, 2.5 or 3 M, prepared by diluting an 8 M urea stock in HEPES buffer. KRysin digestion, quenching and Evotip loading were then carried out as described for HeLa digestion. Each urea concentration was digested in triplicate.

***Automated digestion for phosphoproteomics* (*Figure 3; Supplementary Figures S3 to S8)*** HeLa lysate (400 µg) was digested overnight on a KingFisher Flex robot [24]. Acetonitrile (ACN) was added to the lysates to a concentration of 70% (v/v), and SiMAG-Silanol beads magnetic beads were added at a bead-to-protein ratio of 2:1. Aggregated proteins on bead were washed three times in 500 µL 95% ACN and twice in 70% ethanol. Digestion was performed in 300 µL of 50 mM HEPES pH 8.5, supplemented with 5 mM DTT for KRysin only. Enzymes were added at an enzyme-to-protein ratio of 1:50 or 1:150. For Trypsin/LysC, twice as much Trypsin as LysC was used, giving individual ratios of 1:75 and 1:150 for Trypsin and LysC respectively at a total ratio of 1:50, <u>and similarly to reach a total ratio of 1:150</u>. Digestions were carried out overnight at 37 °C and quenched by addition of formic acid to final 1% (v/v). Six replicates were performed per enzyme and ratio. The equivalent of 500 ng of peptides, based on a digestion recovery of 25%, was loaded directly on Evotips for proteome analysis.

#### Plasma digestion for the sample preparation comparison (Figure 4E; Supplementary Figure S10 to S13)

Three sample preparation strategies were compared: (i) direct digestion, in which 1 µL PPP (approximately 70 µg protein) was diluted in 99 µL 50 mM HEPES pH 8.5, with prior reduction and alkylation using TCEP and chloroacetamide for Trypsin/LysC samples only and 10 mM DTT for KRysin samples only; (ii) PAC [22], in which 1 µL PPP was added to 3 µL 4% SDS, reduced and alkylated with TCEP and chloroacetamide, aggregated onto SiMAG-Silanol magnetic beads by addition of acetonitrile as described above, and digested on-bead in 100 µL 50 mM HEPES pH 8.5 containing 10 mM DTT for KRysin samples; and (iii) urea denaturation, in which 1 µL PPP was added to 9 µL 8 M urea, reduced and alkylated as above (alkylation was not performed for samples to be digested using KRysin), and diluted to 100 µL with 50 mM HEPES pH 8.5 before digestion. A protease-to-plasma-protein ratio of 1:25 was used for all conditions (Trypsin 1:50 plus LysC 1:50, or KRysin 1:25). Digestions were performed at 37 °C with agitation (1000 rpm) and quenched by acidification with a final concentration of 1% TFA (v/v) after 1, 2 and 18 h. Four replicates were performed per condition. An aliquot corresponding to 500 ng was loaded onto Evotips.

#### Plasma digestion for DDA and DIA benchmarking and the LC throughput series (Figure 4B to 4D; Supplementary Figure S9)

Benchmarking of plasma samples for comparison between DDA and DIA on the Orbitrap Exploris 480 mass spectrometer were prepared with urea denaturation as described above. Plasma samples were prepared similarly for experiments evaluating LC throughput.

### Phosphopeptide enrichment

Peptides remaining after the proteome aliquot was taken were desalted on C18 Sep-Pak 96-well plates (40 mg sorbent, Waters) and eluted directly into a KingFisher plate using 75 µL 80% ACN. Phosphopeptides were enriched on a KingFisher Flex robot [24,25] using Zr-IMAC HP magnetic beads (ReSyn Biosciences) at a peptide-to-bead ratio of 1:2. Desalted peptides were supplemented with 200 µL of an adjusted loading buffer to compensate for the elution volume from the desalting step (final 80% ACN, 8% TFA, 0.6 M glycolic acid). Enrichment was carried out as previously described [25], and the enriched phosphopeptides were loaded on Evotips.

### LC-MS/MS data acquisition

#### Liquid chromatography

Peptides were loaded on Evotips (Evosep) according to the manufacturer’s instructions and separated on an Evosep One or Evosep Eno LC system [26]. Unless otherwise noted, the predefined 60, 100, 200, 300, 500 samples-per-day (SPD) methods corresponding to 22.4, 13, 6.4, 4.0, 2.4 minutes per sample, respectively, were used. Phosphopeptides were separated on an Evosep column (8 cm × 150 µm, 1.5 µm, EV1109) connected to a stainless-steel emitter (30 µm i.d., EV1086) at 40 °C using the Evosep Eno 60 SPD method. For the LC throughput series (Figure 4C), the predefined Evosep Eno 60, 100, 200, 300 and 500 SPD methods were used, and peptides were separated on a 4 cm column (4 cm x 150 µm, 1.9 µm, EV1182) at 40 °C.

#### DDA on the Orbitrap Exploris 480 (Figures 1D, 2 and 4B; Supplementary Figures S2, S9 and S14)

An Orbitrap Exploris 480 mass spectrometer (Thermo Fisher Scientific) [27] was operated in positive-ion DDA mode. Full MS1 scans were acquired at a resolution of 60,000, followed by a Top20 method in which the 20 most intense precursor ions per cycle were isolated for higher-energy collisional dissociation (HCD) and MS2 analysis at a resolution of 15,000. Precursor ions were filtered by charge state (2-6), and a dynamic exclusion of 30 s was applied. The MS1 scan range was 350-1400 m/z, with a 1.3 m/z isolation window and a normalized collision energy of 30%.

#### DIA on the Orbitrap Exploris 480 (Figure 4B)

DIA analysis was performed with the following parameters. MS1 resolution of 120,000 and scan range of 350-1400; 49 DIA windows with a width of 13.7 m/z were used. 15,000 MS2 resolution; normalized collision energy % of 35.

#### Targeted DIA of BSA peptides on the Orbitrap Astral (Figure 1B and 1C; Supplementary Figure S1)

An Orbitrap Astral mass spectrometer (Thermo Fisher Scientific) [28] was operated in positive-ion DIA mode. Full MS1 scans were acquired in the Orbitrap analyzer at a resolution of 240,000 across 350 to 1600 m/z. HCD fragmentation was performed in 125 isolation windows (10 m/z wide with 0.5 m/z overlap between adjacent windows) spanning 350 to 1600 m/z, with MS2 spectra acquired in the Astral analyzer at a normalized collision energy of 25%.

#### nDIA of HeLa proteomes and plasma on the Orbitrap Astral (Figures 4C and 4E; Supplementary Figures S3, S4, S10 to S13)

Proteome samples were analyzed on an Orbitrap Astral mass spectrometer using narrow-window data-independent acquisition (nDIA) [20] in positive mode, with a source voltage of 2000 V and an ion transfer tube temperature of 275 °C. MS1 scans were acquired at a resolution of 240,000 over 380 to 980 m/z with an AGC target of 500% and a maximum injection time of 2.5 ms. MS2 spectra were acquired in the Astral analyzer over 150 to 2000 m/z after HCD at a normalized collision energy of 25%, using 299 DIA windows with 2 m/z isolation width per window, a maximum injection time of 2.5 ms and an AGC target of 500%. For plasma samples analyzed using different SPD methods as presented in Figure 4C, maximum injection time of 1.5 ms, and AGC target of 200% was used.

### nDIA of phosphopeptides on the Orbitrap Astral (Figure 3; Supplementary Figures S5 to S8)

Enriched phosphopeptides were analyzed with the same nDIA settings, except that MS1 scans covered 480 to 1080 m/z with a maximum injection time set at 3 ms. Tandem mass spectra were generated over 150 to 2,000 m/z after HCD fragmentation with a normalized collision energy of 25%. 2 Th isolation windows were used with a maximum injection time of 2.5 ms and an AGC target of 500%.

### Data processing

Raw data were processed separately from acquisition as described below.

#### Sequence databases

Human data were searched against the UniProt human reference proteome (UniProt reference proteome 2023 release, 20,598 entries) alongside a reference proteome database of standard proteomic contaminants. BSA data were analyzed using the UniProt entry P02769.

#### DDA data processing in MaxQuant (Figures 2 and 4B; Supplementary Figures S2, S9 and S14)

DDA raw files were processed using MaxQuant 2.6.7.0 [17,29] with enzyme specificity set to Trypsin/P or Trypsin/P plus LysC/P and up to 3 missed cleavages. Carbamidomethylation of cysteine was set as a fixed modification, and protein N-terminal acetylation and methionine oxidation as variable modifications. Minimum peptide length of 7 amino acids with 1% FDR thresholds at PSM and protein level was used. Match between runs was set to off.

#### Unspecific search for C-terminal specificity (Figure 1D; Supplementary Table S1)

To determine C-terminal cleavage specificity without assuming protease rules, HeLa DDA raw files from KRysin and Trypsin digests were searched in MaxQuant [17] with no protease specificity defined and digestion mode set to “unspecific”. 1% FDR was used at the PSM and protein group level. The C-terminal residue of each identified peptide was extracted from the peptide output, giving 8,755 KRysin and 8,316 Trypsin peptides.

#### Targeted analysis of BSA peptides in Skyline (Figure 1B and 1C; Supplementary Figure S1)

Targeted extraction of BSA peptides from the Orbitrap Astral DIA data was performed in Skyline 26.1.0 [30]. The peptide list was generated from the BSA sequence (UniProt P02769), specifying full cleavage C-terminal to lysine and arginine with no missed cleavage, no cleavage exception at proline and no cysteine modification. Precursor and fragment ion chromatograms were extracted from the DIA isolation windows based on theoretical precursor m/z values, and peak identification was confirmed based on retention time consistency, precursor mass accuracy and fragment ion co-elution. Precursor charge states of 2 and 3 were used, singly charged b and y fragment ion were monitored. Mass tolerance of 0.5 m/z, minimum peptide length of 6 amino acid was used to define the 43 predicted BSA peptides. Extracted peptide chromatograms were manually inspected to ensure correct peak integration.

#### nDIA data processing in Spectronaut (Figure 3; Supplementary Figures S3 to S8)

nDIA raw files for data presented in Figure 3 were searched library free using Spectronaut (v21) [31] against the human protein reference database, supplemented with a contaminant database from Spectronaut. Cysteine carbamidomethylation was set as a fixed modification, and N-terminal acetylation and methionine oxidation were set as variable modifications, with a maximum number of variable modifications kept at 5, and a maximum number of missed cleavages kept at 2. The maximum and minimum number of fragments per peptide were set to 25 and 6, respectively. Cross-run normalisation was not selected, and method evaluation was allowed, where conditions were defined per enzyme and per ratio. For phosphoproteomics analysis, serine, threonine, and tyrosine phosphorylation were added as variable modifications, and the PTM localization filter was activated with a threshold of 0.75 (class I sites) [32].

#### Plasma LC throughput data processing in DIA-NN (Figures 4C to 4E; Supplementary Figures S10 to S13)

Plasma nDIA and Orbitrap Exploris 480 data were processed with DIA-NN version 2.0.2 [33]. The data were searched in library-free mode against the UniProt human reference proteome (2023 release, 20,598 entries) with Trypsin as the cleavage enzyme and 1 missed cleavage allowed. FDR was set to 1%. Cysteine carbamidomethylation was set as a fixed modification for alkylated samples (Trypsin, Trypsin/LysC and the PAC-prepared KRysin samples) and was omitted for KRysin samples digested without alkylation (direct and urea workflows).

### Data analysis

#### Missed-cleavage and cleavage-efficiency metrics

Peptide-level missed-cleavage rates were calculated as the percentage of unique identified peptide sequences with 0, 1, 2 or 3 internal lysine or arginine residues (Figures 2B to 2D, 2F and 3D, and Supplementary Figures S3, S9A and S9B, S10 and S11).

#### Phosphoproteomics analysis (Figure 3; Supplementary Figures S3 to S8)

Phosphopeptides and class I phosphosites were counted per sample and as identifications valid in all six replicates or in four or five of six replicates (Figure 3B and 3C). Phosphosites were defined by their gene, modification site and multiplicity, and multiplicity was calculated as the number of phosphorylated residues per phosphopeptide (Figure 3F). The overlap between Trypsin/LysC and KRysin at 1:50 was based on sites quantified in at least four of six replicates in both Trypsin/LysC and KRysin conditions (Supplementary Figure S6). For sites unique to either enzyme or shared, the supporting phosphopeptides were characterized by missed-cleavage status, peptide length, the distance between adjacent lysine or arginine residues and hydrophobicity [34] (Supplementary Figure S7). Sequence motifs of KRysin-unique sites relative to Trypsin/LysC-unique sites were analyzed with iceLogo [35] using a window of seven residues on either side of the phosphosite, percentage difference scoring and P < 0.1 (Supplementary Figure S8).

For the salt-bridge analysis (Figure 3G), salt bridges were defined by the presence of a lysine or arginine at position −2 or +2 relative to the phosphorylated site. A missed cleavage was attributed to the salt-bridge position when the amino acids from position −2 to the modified site in the flanking region matched the first three amino acids of the peptide sequence (lysine or arginine at −2), or when the amino acids from the modified site to position +2 matched the last three amino acids of the peptide sequence. Only missed cleavages from peptides bearing a phosphosite in a salt-bridge position to which the missed cleavage could be attributed were considered. The share of intensity from fully cleaved and miscleaved phosphopeptides was then calculated for these sites.

#### Plasma protein group analysis (Figure 4C and 4D)

Protein groups were counted per replicate, and medians of triplicate digestions are reported (Figure 4C). KRysin-specific protein groups were defined as protein groups identified in all three KRysin replicates at any LC throughput and not identified with Trypsin or Trypsin/LysC at any throughput. Protein groups were assigned to Gene Ontology (GO) Biological Process, Molecular Function and Cellular Component terms using g:Profiler g:GOSt [36] with GO annotations from Ensembl BioMart (release 2026-01-23). The 473 KRysin-specific protein groups (387 of which mapped to a recognized accession in the g:Profiler namespace) were tested against a custom statistical background comprising the union of all protein groups detected with any enzyme at any LC throughput, mapped to 1,102 UniProt accessions (effective annotated domain size, 1,254 genes), using Fisher’s exact test with Benjamini-Hochberg FDR correction. Terms with q < 0.05 were considered significant.

#### Quantitative reproducibility (Supplementary Figure S12 and S13)

Protein group intensities from DIA-NN, exported without cross-run normalization, were used to assess quantitative reproducibility across the four replicate digests of each enzyme, sample preparation method and digestion time. After removal of contaminants, coefficients of variation (CVs) were calculated for each protein group quantified in at least three of four replicates as the standard deviation divided by the mean intensity. To separate protein-level variation from differences in total signal between replicate digests, CVs were also calculated after per-run median normalization. Each run was scaled by the median ratio of its intensities to the per-protein median across replicates, using protein groups quantified in all four replicates. Differences in total signal were assessed from the summed protein group intensity of each replicate digest, expressed relative to the condition mean (Supplementary Figure S12). To assess reproducibility as a function of abundance, unnormalized CVs were plotted against log2 mean protein intensity and fitted with LOESS trend lines with 95% confidence intervals, with the y-axis limited to the 95th percentile of all CVs (Supplementary Figure S13). For summary statistics, protein groups in each condition were divided into thirds by mean intensity.

#### Statistics and software

Unless otherwise stated, replicates are independent digestions of the same lysate or plasma sample, bars show means and error bars show standard deviations. Data analysis and visualization were performed in R version 4.4.1 [37] using RStudio 2024.12.1 as the integrated development environment.

## Supplementary Information

Supplementary Figures S1 to S14 and Supplementary Tables S1 and S2 are provided in the Supplementary Information file.

## Author Contributions

T.S.B.: Conceptualization, methodology, investigation, formal analysis, data curation, visualization, writing, review and editing. C.H.R.: Conceptualization, methodology, investigation, formal analysis, data curation, writing, review and editing. C.K.: Investigation, formal analysis, data curation, writing, review and editing. F.L.M.: review and editing. J.V.O.: Conceptualization, supervision, funding acquisition, project administration, review and editing.

## Data Availability

Raw MS data and search outputs will be deposited to ProteomeXchange via PRIDE upon submission; accession number to be added.

## Acknowledgements

Work at The Novo Nordisk Foundation Center for Protein Research (CPR) is funded in part by a donation from the Novo Nordisk Foundation (NNF21OC0072070, and NNF24SA0098829). This project was supported by a center-of-excellence grant from the Danish National Research Foundation to Copenhagen Center for Glycocalyx Research (DNRF196). The work was also supported by PLATO, a grant from the Danish Ministry of Higher Education and Science to establish the PLATO research infrastructure: Danish National Mass Spectrometry Platform for Proteomics and Biomolecular Imaging (grant no. 5229-00012B, www.sdu.dk/PLATO ).

