## Supplementary information for "High-fidelity bispecific protease KRysin for deeper plasma proteomics and phosphoproteomics"

### Supplementary Figure S1: Extracted Ion Chromatograms for KRysin BSA digested “tryptic” peptides (first page)

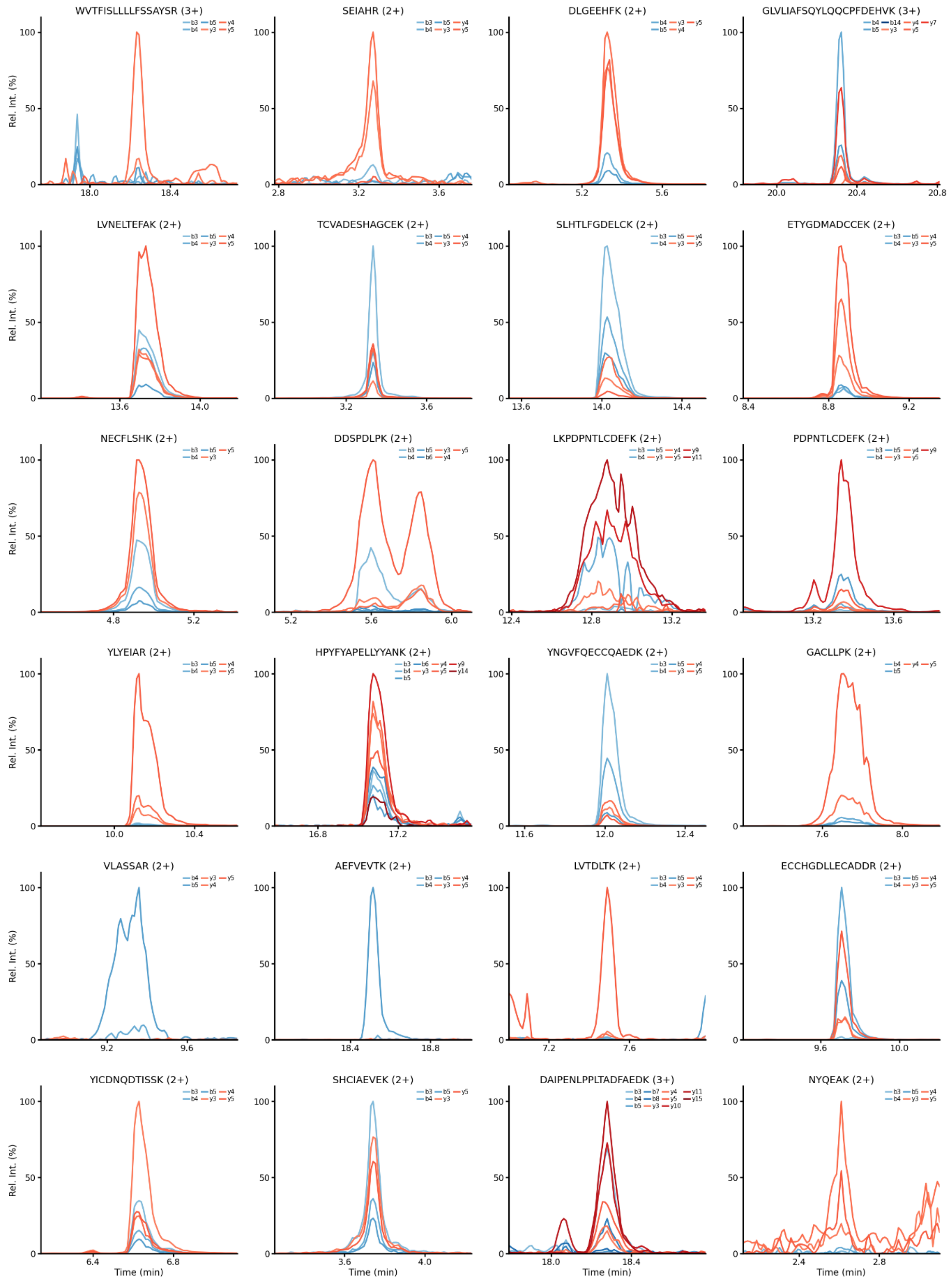

Continued on next page

### Supplementary Figure 1: Extracted Ion Chromatograms for KRysin BSA digested “tryptic” peptides (second page)

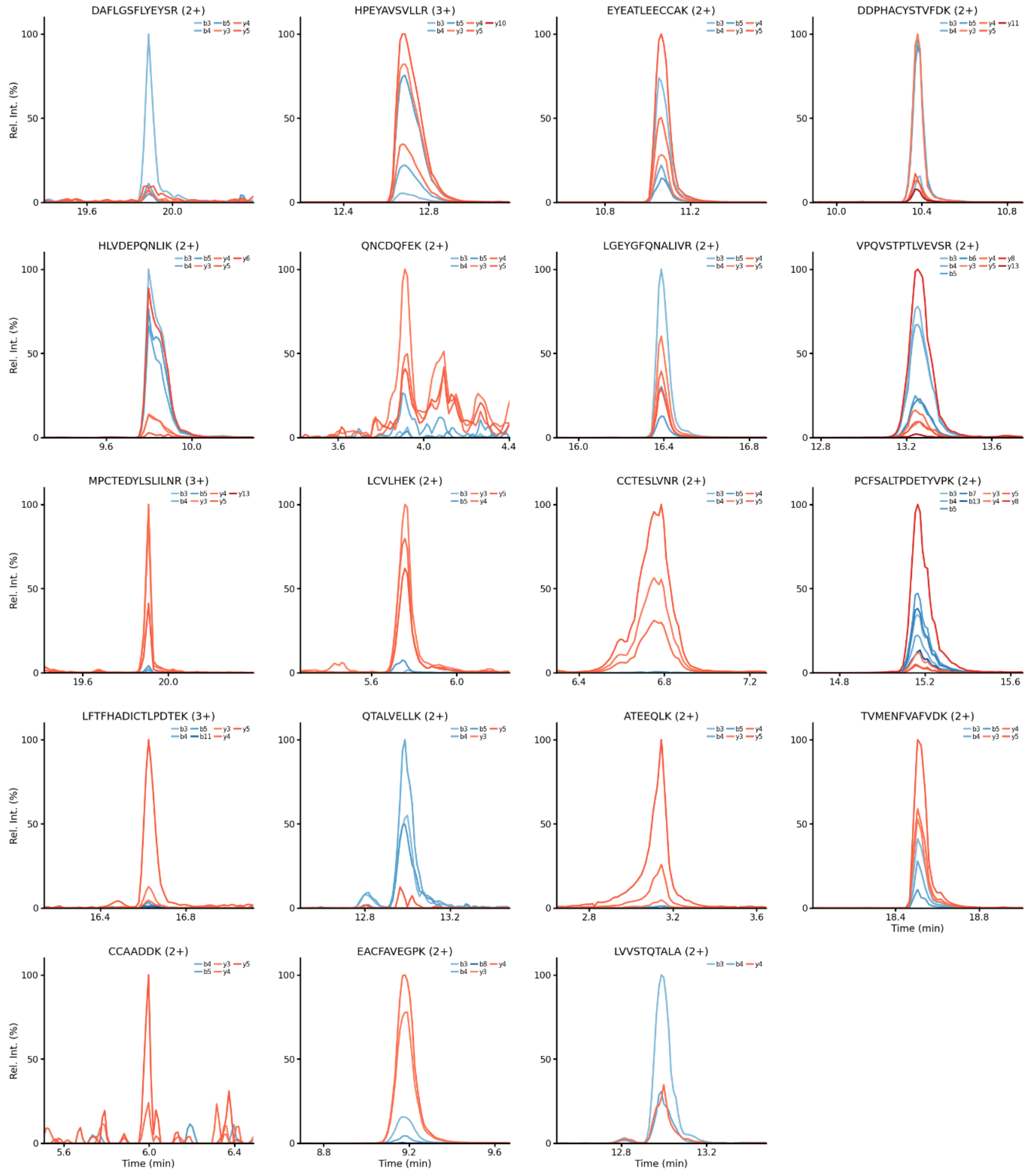

Extracted ion chromatograms (XICs) are shown for the 43 tryptic BSA peptides monitored in this targeted proteomics experiment, distributed across two pages. Each panel displays the precursor sequence and charge state as the title, with retention time (min) on the x-axis and relative intensity (%) on the y-axis, normalized to the most intense product ion trace within the displayed elution window for that peptide. Each trace corresponds to a single product ion (fragment), color-coded by ion series and rank: b-ions are shown in blue shades and y-ions in red shades, with darker shades indicating higher-numbered ions within each series; the specific fragment ions monitored for each peptide are indicated in the panel legend. Chromatographic windows were centered on the apex of the summed product ion signal for each peptide and cropped to a 1-minute window (±0.5 min) to allow direct visual comparison of peak shape and co-elution across product ions.

#### Supplementary Table S1. C-terminal amino acid specificity of KRysin and Trypsin determined by unspecific search analysis

| Last amino acid | Count KRysin | Count Trypsin | % KRysin | % Trypsin |
|---|---|---|---|---|
| R | 3683 | 3416 | 42,07% | 41,08% |
| K | 4979 | 4720 | 56,87% | 56,76% |
| E | 8 | 15 | 0,09% | 0,18% |
| V | 6 | 9 | 0,07% | 0,11% |
| H | 3 | 17 | 0,03% | 0,20% |
| Q | 11 | 4 | 0,13% | 0,05% |
| L | 7 | 22 | 0,08% | 0,26% |
| Y | 4 | 10 | 0,05% | 0,12% |
| D | 8 | 16 | 0,09% | 0,19% |
| N | 8 | 20 | 0,09% | 0,24% |
| F | 5 | 17 | 0,06% | 0,20% |
| M | 2 | 7 | 0,02% | 0,08% |
| A | 13 | 19 | 0,15% | 0,23% |
| I | 5 | 7 | 0,06% | 0,08% |
| G | 4 | 3 | 0,05% | 0,04% |
| S | 5 | 8 | 0,06% | 0,10% |
| T | 4 | 3 | 0,05% | 0,04% |
| C | 0 | 3 | 0,00% | 0,04% |
| Total | 8755 | 8316 | | |

Distribution of C-terminal amino acid residues among peptides identified from KRysin and Trypsin digests of HeLa cell protein extracts, determined by unspecific search analysis of LC-MS/MS data. For each protease, peptides were grouped by their C-terminal residue, and the count and percentage of total identified peptides are reported. Both proteases show the expected predominance of C-terminal lysine and arginine, consistent with K/R-specific cleavage specificity, with a minor fraction of peptides terminating in other residues reflecting non-specific cleavage. Total peptide counts were 8,755 for KRysin and 8,316 for Trypsin.

**Supplementary Figure S2. Residue-specific missed cleavage rates for Arginine and Lysine following KRysin, Trypsin, and Trypsin/LysC digestion of a HeLa protein extract.**

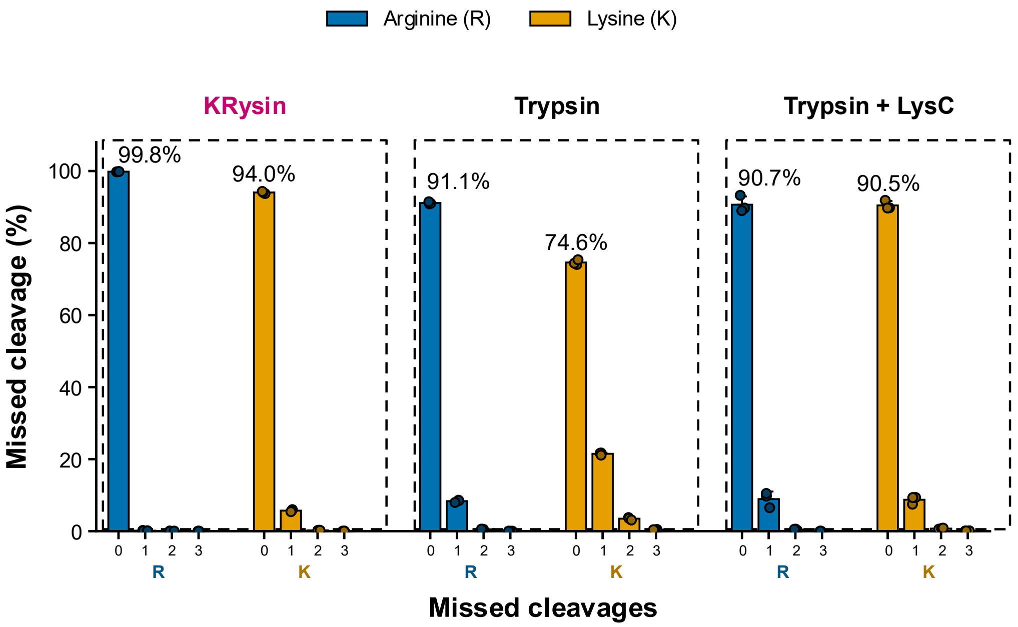

Percentage of peptides with 0 to 3 missed cleavages at arginine (R) or lysine (K), determined separately for each residue type following digestion of a HeLa cell lysate protein extract with KRysin, Trypsin, or Trypsin/LysC. Bars represent the mean of three technical replicates, error bars indicate the standard deviation, and individual points show the value for each replicate. The percentage of fully cleaved (0 missed cleavage) sites is annotated above the corresponding bar.

#### Supplementary Table S2. Peptide counts and percentage of peptides at each missed cleavage level across urea concentrations for KRysin digestion of a HeLa protein extract.

Peptide counts (A) and the corresponding percentage of peptides (B) at each missed cleavage level (0 to 3) following KRysin digestion of a HeLa cell lysate protein extract across a urea concentration gradient (0 to 3 M). Values are shown for each of three technical replicates (Rep 1–Rep 3) together with the mean ± standard deviation (SD).

##### A. Peptide count

| Urea conc. | Missed cleavages | Rep 1 | Rep 2 | Rep 3 | Mean ± SD |
|---|---|---|---|---|---|
| **0M** | 0 | 11,068 | 10,854 | 10,789 | **10,904 ± 146** |
| | 1 | 637 | 578 | 560 | **592 ± 40** |
| | 2 | 25 | 26 | 21 | **24 ± 3** |
| | 3 | 1 | 2 | 0 | **1 ± 1** |
| **0.5M** | 0 | 10,909 | 11,107 | 10,737 | **10,918 ± 185** |
| | 1 | 550 | 575 | 520 | **548 ± 28** |
| | 2 | 21 | 22 | 17 | **20 ± 3** |
| | 3 | 1 | 0 | 0 | **0 ± 1** |
| **1M** | 0 | 9,592 | 10,168 | 9,889 | **9,883 ± 288** |
| | 1 | 637 | 740 | 686 | **688 ± 52** |
| | 2 | 32 | 40 | 40 | **37 ± 5** |
| | 3 | 2 | 5 | 1 | **3 ± 2** |
| **1.5M** | 0 | 8,830 | 8,531 | 9,763 | **9,041 ± 643** |
| | 1 | 604 | 634 | 647 | **628 ± 22** |
| | 2 | 27 | 28 | 32 | **29 ± 3** |
| | 3 | 3 | 3 | 0 | **2 ± 2** |
| **2M** | 0 | 7,350 | 8,875 | 8,656 | **8,294 ± 825** |
| | 1 | 928 | 911 | 864 | **901 ± 33** |
| | 2 | 68 | 67 | 51 | **62 ± 10** |
| | 3 | 9 | 6 | 4 | **6 ± 3** |
| **2.5M** | 0 | 5,668 | 9,263 | 9,104 | **8,012 ± 2,031** |
| | 1 | 310 | 744 | 900 | **651 ± 306** |
| | 2 | 11 | 34 | 61 | **35 ± 25** |
| | 3 | 0 | 3 | 5 | **3 ± 3** |
| **3M** | 0 | 5,747 | 6,430 | 6,498 | **6,225 ± 415** |
| | 1 | 1,321 | 1,237 | 1,009 | **1,189 ± 161** |
| | 2 | 180 | 140 | 85 | **135 ± 48** |
| | 3 | 23 | 17 | 8 | **16 ± 8** |

#### B. Missed cleavage (%)

| Urea conc. | Missed cleavages | Rep 1 | Rep 2 | Rep 3 | Mean ± SD |
|---|---|---|---|---|---|
| **0M** | 0 | 94.35 | 94.71 | 94.89 | **94.65 ± 0.28** |
| | 1 | 5.43 | 5.04 | 4.93 | **5.13 ± 0.26** |
| | 2 | 0.21 | 0.23 | 0.18 | **0.21 ± 0.02** |
| | 3 | 0.01 | 0.02 | 0.00 | **0.01 ± 0.01** |
| **0.5M** | 0 | 95.02 | 94.90 | 95.24 | **95.05 ± 0.17** |
| | 1 | 4.79 | 4.91 | 4.61 | **4.77 ± 0.15** |
| | 2 | 0.18 | 0.19 | 0.15 | **0.17 ± 0.02** |
| | 3 | 0.01 | 0.00 | 0.00 | **0.00 ± 0.01** |
| **1M** | 0 | 93.46 | 92.83 | 93.15 | **93.15 ± 0.31** |
| | 1 | 6.21 | 6.76 | 6.46 | **6.47 ± 0.27** |
| | 2 | 0.31 | 0.37 | 0.38 | **0.35 ± 0.03** |
| | 3 | 0.02 | 0.05 | 0.01 | **0.02 ± 0.02** |
| **1.5M** | 0 | 93.30 | 92.77 | 93.50 | **93.19 ± 0.38** |
| | 1 | 6.38 | 6.89 | 6.20 | **6.49 ± 0.36** |
| | 2 | 0.29 | 0.30 | 0.31 | **0.30 ± 0.01** |
| | 3 | 0.03 | 0.03 | 0.00 | **0.02 ± 0.02** |
| **2M** | 0 | 87.97 | 90.02 | 90.40 | **89.46 ± 1.31** |
| | 1 | 11.11 | 9.24 | 9.02 | **9.79 ± 1.15** |
| | 2 | 0.81 | 0.68 | 0.53 | **0.68 ± 0.14** |
| | 3 | 0.11 | 0.06 | 0.04 | **0.07 ± 0.03** |
| **2.5M** | 0 | 94.64 | 92.22 | 90.41 | **92.42 ± 2.12** |
| | 1 | 5.18 | 7.41 | 8.94 | **7.17 ± 1.89** |
| | 2 | 0.18 | 0.34 | 0.61 | **0.38 ± 0.21** |
| | 3 | 0.00 | 0.03 | 0.05 | **0.03 ± 0.02** |
| **3M** | 0 | 79.04 | 82.18 | 85.50 | **82.24 ± 3.23** |
| | 1 | 18.17 | 15.81 | 13.28 | **15.75 ± 2.45** |
| | 2 | 2.48 | 1.79 | 1.12 | **1.79 ± 0.68** |
| | 3 | 0.32 | 0.22 | 0.11 | **0.21 ± 0.11** |

**Supplementary Figure S3. Missed cleavage rates of non-phosphorylated peptides by peptide count.**

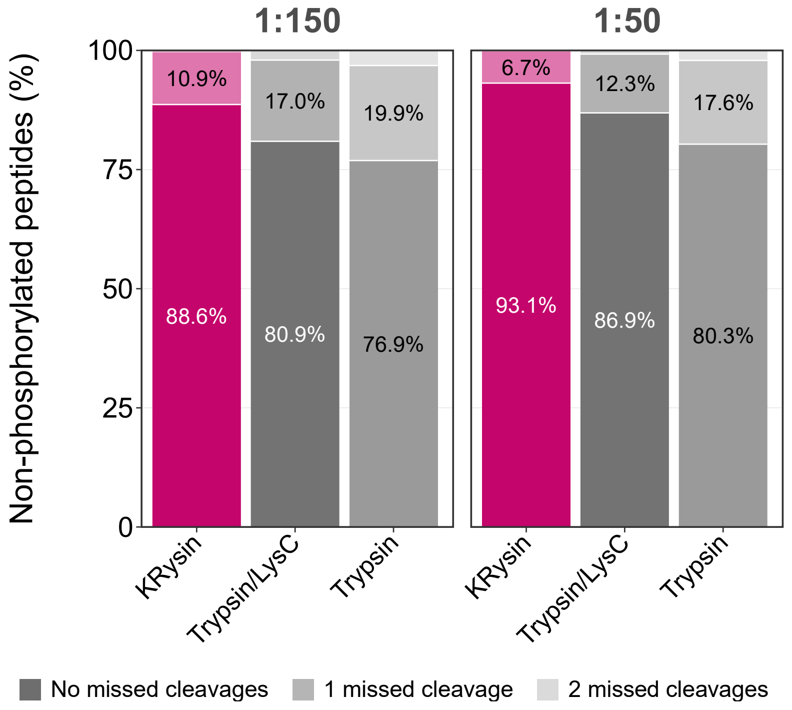

Stacked bar plots show the percentage of non-phosphorylated peptides identified with no missed cleavages, one missed cleavage, or two missed cleavages, calculated from peptide counts. HeLa protein extracts were digested with KRysin, Trypsin/LysC, or Trypsin at enzyme-to-substrate ratios of 1:150 (left) and 1:50 (right), and resulting peptides analyzed by nDIA on an Orbitrap Astral mass spectrometer. At 1:150, peptides with no missed cleavages accounted for 88.6% of KRysin, 80.9% of Trypsin/LysC, and 76.9% of Trypsin identifications. At 1:50, the corresponding values were 93.1%, 86.9%, and 80.3%. Peptides with one missed cleavage accounted for 10.9%, 17.0%, and 19.9% at 1:150, and 6.7%, 12.3%, and 17.6% at 1:50, for KRysin, Trypsin/LysC, and Trypsin respectively. Mean of n = 6 replicates.

**Supplementary Figure S4. MS2 signal intensities across digestion conditions.**

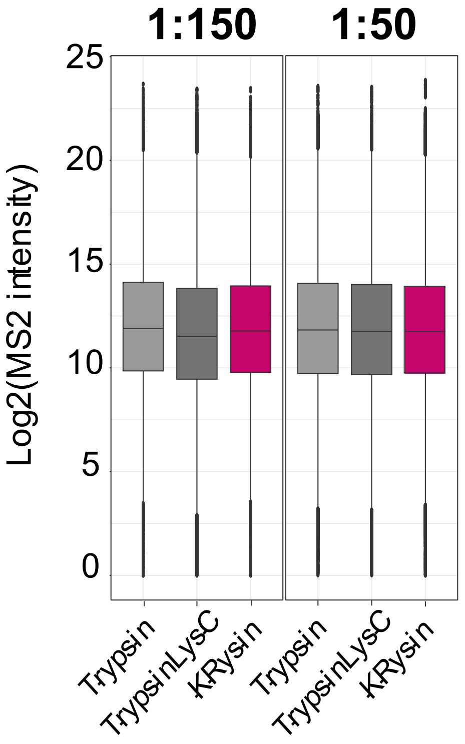

Box plots show the distribution of log2-transformed MS2 intensities for peptides identified from HeLa protein extracts digested with Trypsin, Trypsin/LysC, or KRysin at enzyme-to-substrate ratios of 1:150 (left) and 1:50 (right), analyzed by nDIA on an Orbitrap Astral mass spectrometer. Boxes span the interquartile range, the center line indicates the median, whiskers extend to 1.5 times the interquartile range, and points beyond the whiskers indicate outliers.

#### Supplementary Figure S5. Class 1 phosphosites by phosphorylated residue across digestion conditions.

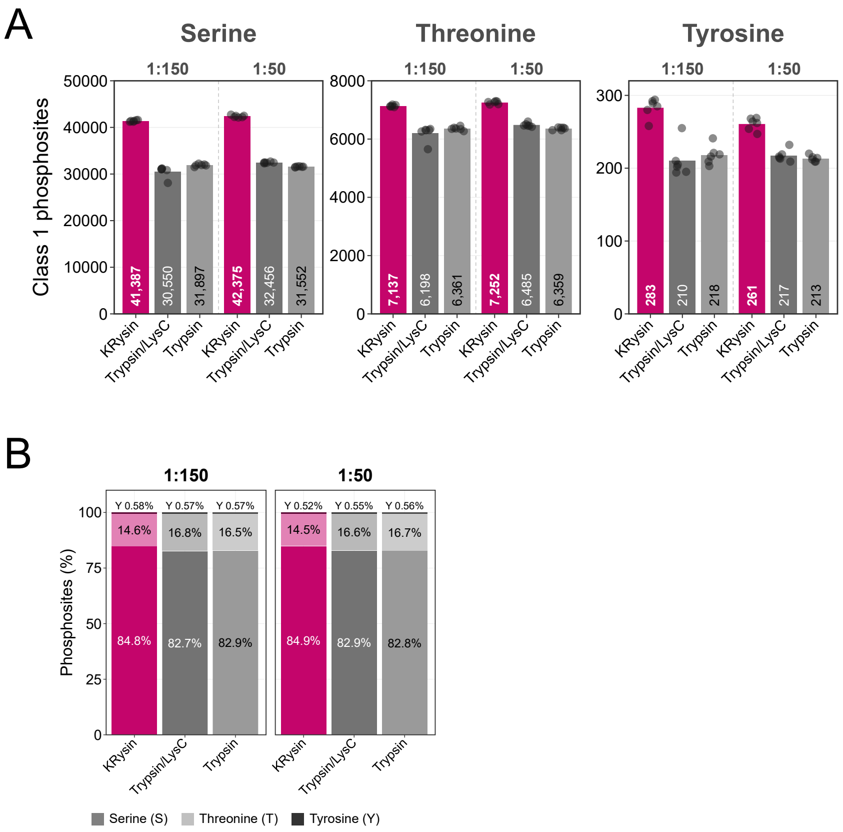

(A) Number of class 1 phosphosites localized to serine, threonine, and tyrosine residues identified from HeLa protein extracts digested with KRysin, Trypsin/LysC, or Trypsin at enzyme-to-substrate ratios of 1:150 and 1:50. Bars indicate the mean and points indicate individual replicates (n = 6). (B) Relative distribution of serine, threonine, and tyrosine among class 1 phosphosites for each digestion condition, expressed as a percentage of total class 1 phosphosites. Samples were analyzed by nDIA on an Orbitrap Astral mass spectrometer.

**Supplementary Figure S6. Overlap of quantified phosphosites between Trypsin/LysC and KRysin digestions and their phosphorylation multiplicity.**

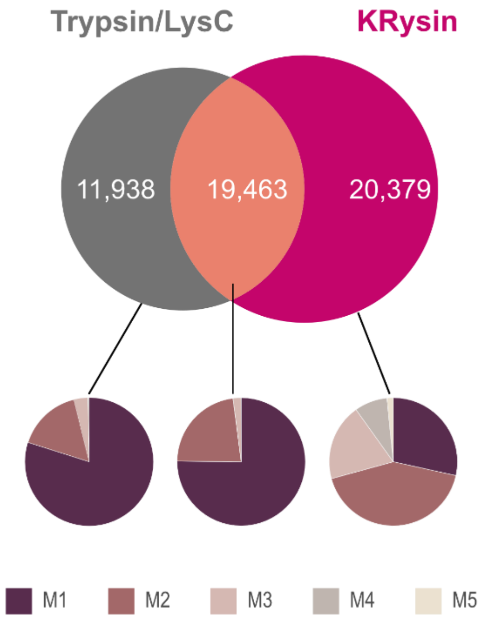

Venn diagram showing the overlap of quantified class 1 phosphosites identified from HeLa protein extracts digested with Trypsin/LysC or KRysin at an enzyme-to-substrate ratio of 1:50 and quantified in at least four of six replicates, analyzed by nDIA on an Orbitrap Astral mass spectrometer. In total, 11,938 sites were unique to Trypsin/LysC, 19,463 were shared between the two conditions, and 20,379 were unique to KRysin. Pie charts below show the distribution of phosphorylation multiplicity (M1 to M5, corresponding to one to five phosphorylated residues on the peptide) for the sites unique to Trypsin/LysC, shared between both conditions, and unique to KRysin, respectively.

**Supplementary Figure S7. Characteristics of phosphopeptides supporting enzyme-unique and shared phosphosites.**

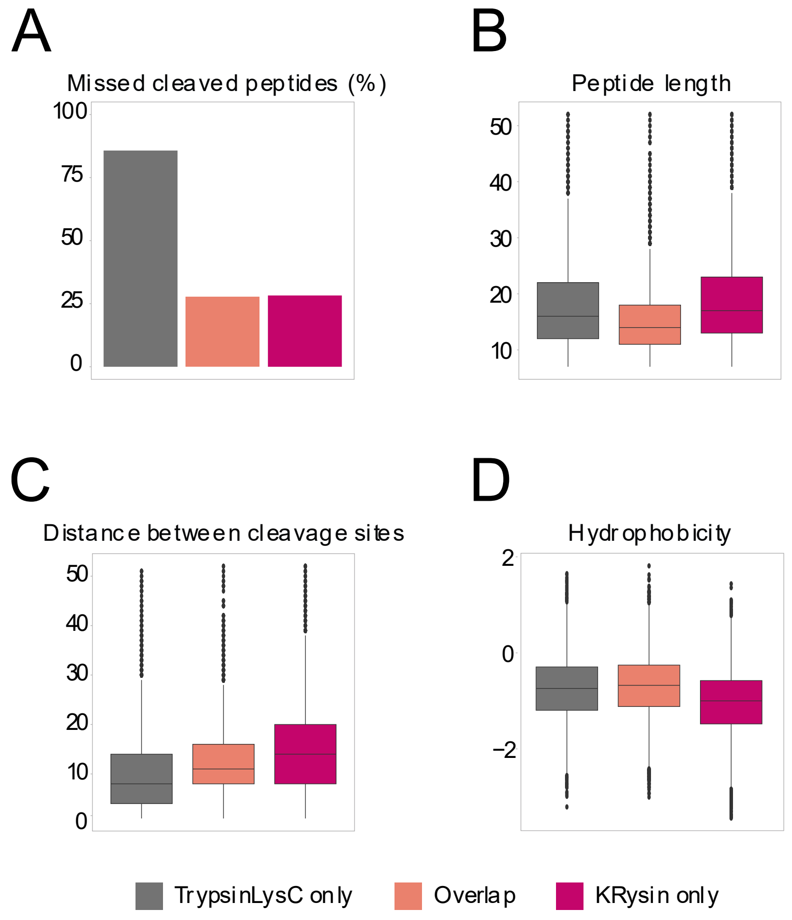

Phosphosites were grouped as unique to Trypsin/LysC, shared between both digestions (overlap), or unique to KRysin, as defined in Supplementary Figure S6. For the phosphopeptides supporting each group, the plots show (**A**) the percentage of peptides containing at least one missed cleavage, (**B**) peptide length in amino acids, (**C**) the distance in amino acids between adjacent cleavage sites (lysine and arginine residues), and (D) peptide hydrophobicity (Kyte-Doolittle GRAVY). In box plots, the center line indicates the median, boxes span the interquartile range, whiskers extend to 1.5 times the interquartile range, and points indicate outliers. HeLa protein extracts were analyzed by nDIA on an Orbitrap Astral mass spectrometer.

**Supplementary Figure S8. Sequence motif analysis of phosphosites unique to KRysin digestion.**

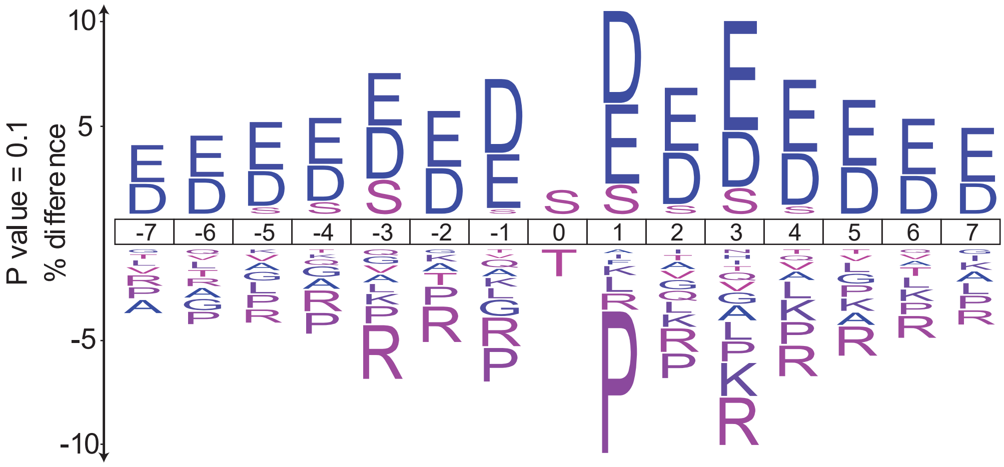

Sequence logo of the amino acids surrounding class 1 phosphosites unique to KRysin relative to those unique to Trypsin/LysC, shown as a window of seven residues on either side of the phosphorylated residue (positions -7 to +7, with the phosphosite at position 0). Residues above the axis are enriched and residues below the axis are depleted in sites unique to KRysin, with letter height indicating the percentage difference in residue frequency between the two sets. Only residues with a significant difference at P < 0.1 are shown. Sites were derived from HeLa protein extracts digested with KRysin or Trypsin/LysC and analyzed by nDIA on an Orbitrap Astral mass spectrometer.

**Supplementary Figure S9. Missed cleavage profiles of peptides from human platelet-poor plasma digested with Trypsin, Trypsin/LysC, or KRysin.**

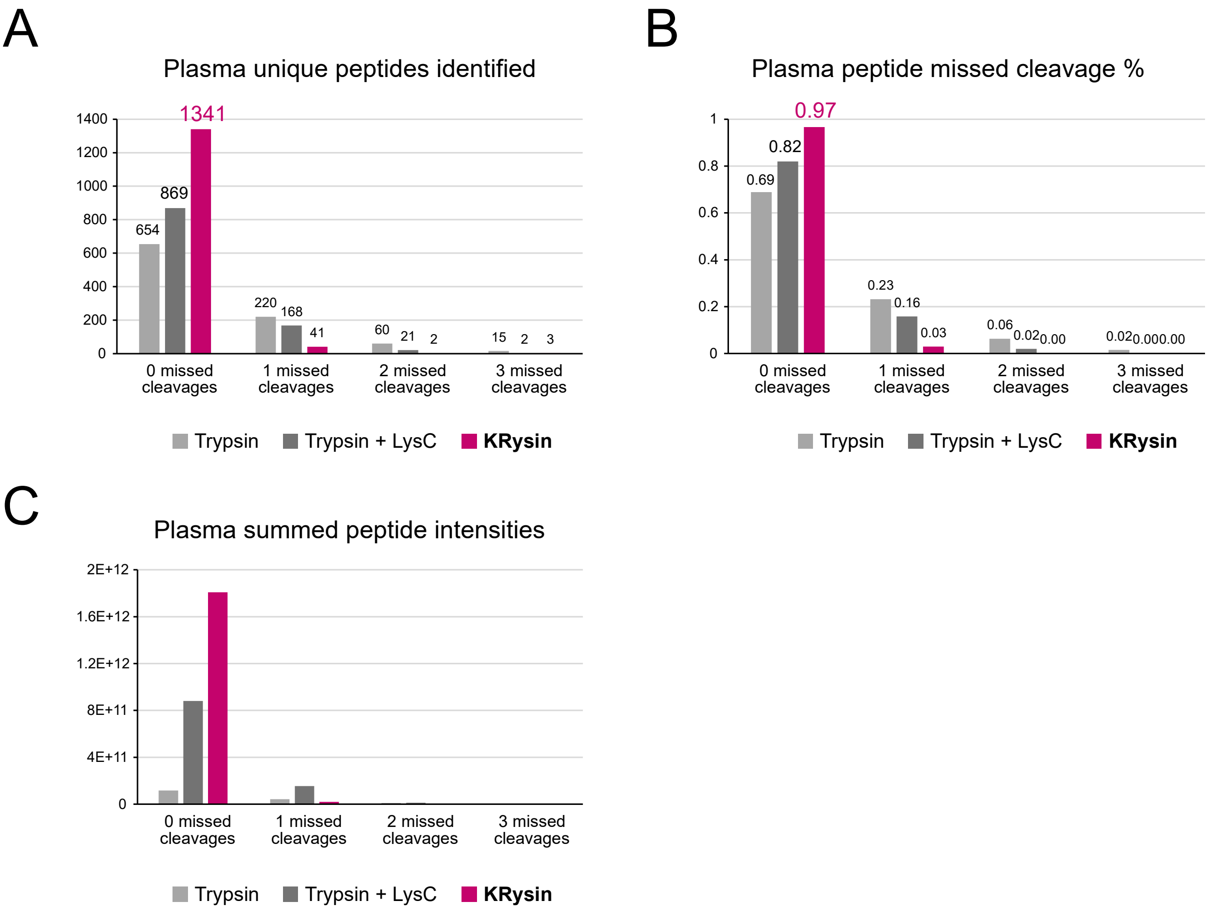

Human platelet-poor plasma was digested with Trypsin, Trypsin/LysC, or KRysin and analyzed by DDA LC-MS/MS on a Thermo Fisher Orbitrap Exploris 480 mass spectrometer. Identified peptides were grouped by the number of missed cleavages (0 to 3). (A) Number of unique peptides identified in each missed cleavage category. KRysin yielded 1341 unique peptides with no missed cleavages, compared with 869 for Trypsin/LysC and 654 for Trypsin, and fewer peptides with one missed cleavage (41, compared with 168 and 220, respectively). (B) Fraction of unique peptides in each missed cleavage category. Peptides without missed cleavages made up 0.97 of the peptides identified with KRysin, compared with 0.82 for Trypsin/LysC and 0.69 for Trypsin. (C) Summed peptide intensities in each missed cleavage category. Summed intensity of peptides without missed cleavages was highest for KRysin, followed by Trypsin/LysC and then Trypsin.

**Supplementary Figure S10. Missed cleavage rates in plasma digests analyzed by nDIA across sample preparation methods and digestion times.**

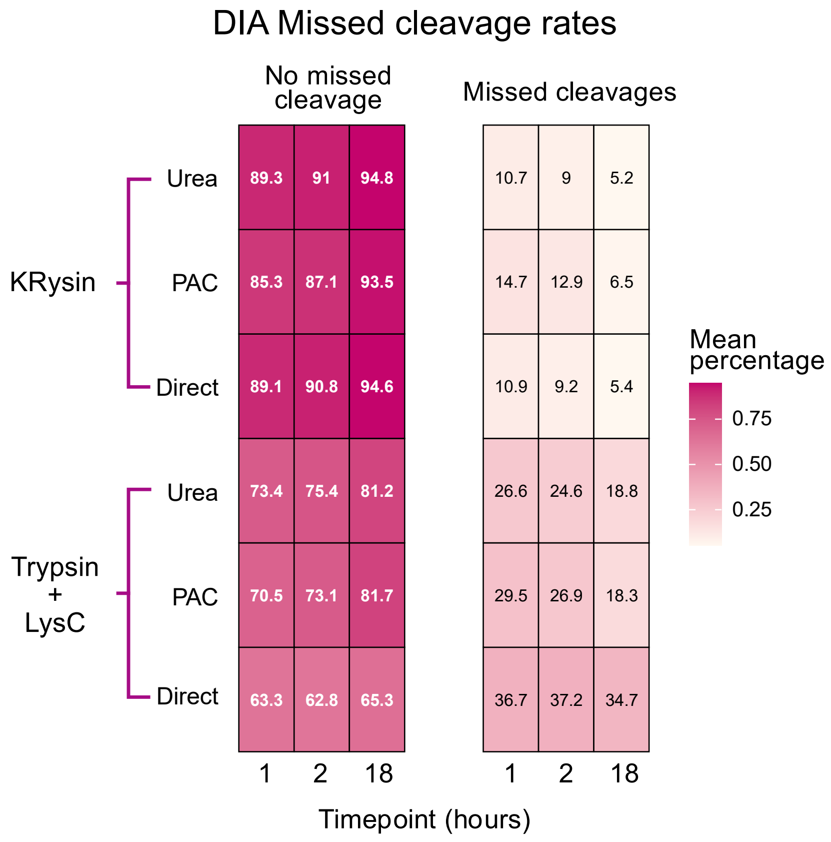

Heatmaps show the mean percentage of identified peptides from human plasma digests with no missed cleavages (left) or with one or more missed cleavages (right), based on peptide counts. Human platelet-poor plasma was digested with KRysin or Trypsin/LysC for 1, 2, or 18 hours using three sample preparation methods: direct digestion in buffer (Direct), protein aggregation capture (PAC), or digestion in the presence of urea (Urea). Protease-to-plasma protein ratio was 1:25 for both enzymes (1:50 each for Trypsin and LysC). Values in each cell are the mean percentage (n = 4 replicates), and color intensity reflects the value. After 1 hour, the mean missed cleavage rate was 10.7 to 14.7% for KRysin and 26.6 to 36.7% for Trypsin/LysC across the three sample preparation methods.

**Supplementary Figure S11. Unique peptide identifications from human platelet-poor plasma digested with KRysin or Trypsin/LysC, separated by missed cleavage status.**

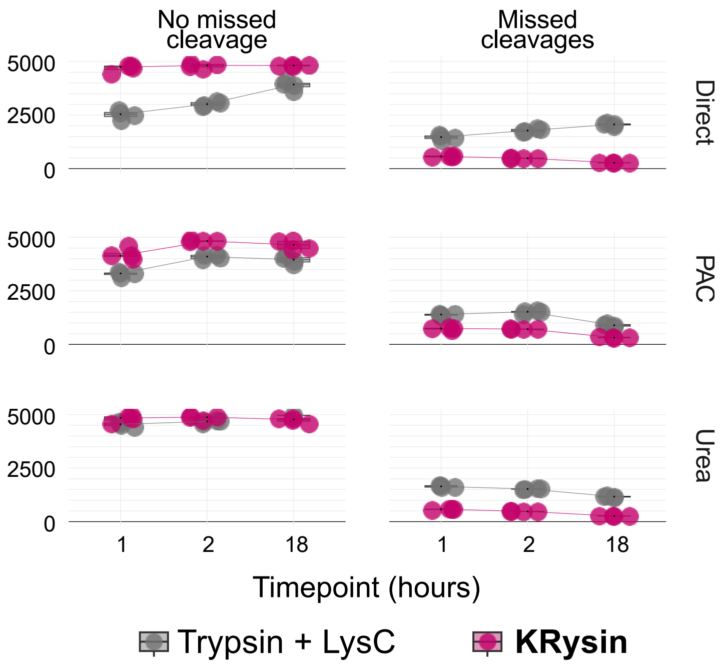

Human platelet-poor plasma was digested with KRysin or Trypsin/LysC for 1, 2, or 18 hours using three sample preparation workflows (Direct, PAC, and Urea) and analyzed by narrow-window data-independent acquisition (DIA) on an Orbitrap Astral mass spectrometer. The number of unique peptides identified is shown for peptides with no missed cleavages (left) and peptides with missed cleavages (right). Points represent individual replicate digests, and lines connect the mean at each timepoint.

#### Supplementary Figure S12. Reproducibility of plasma protein quantification across preparation methods with KRysin and trypsin/LysC.

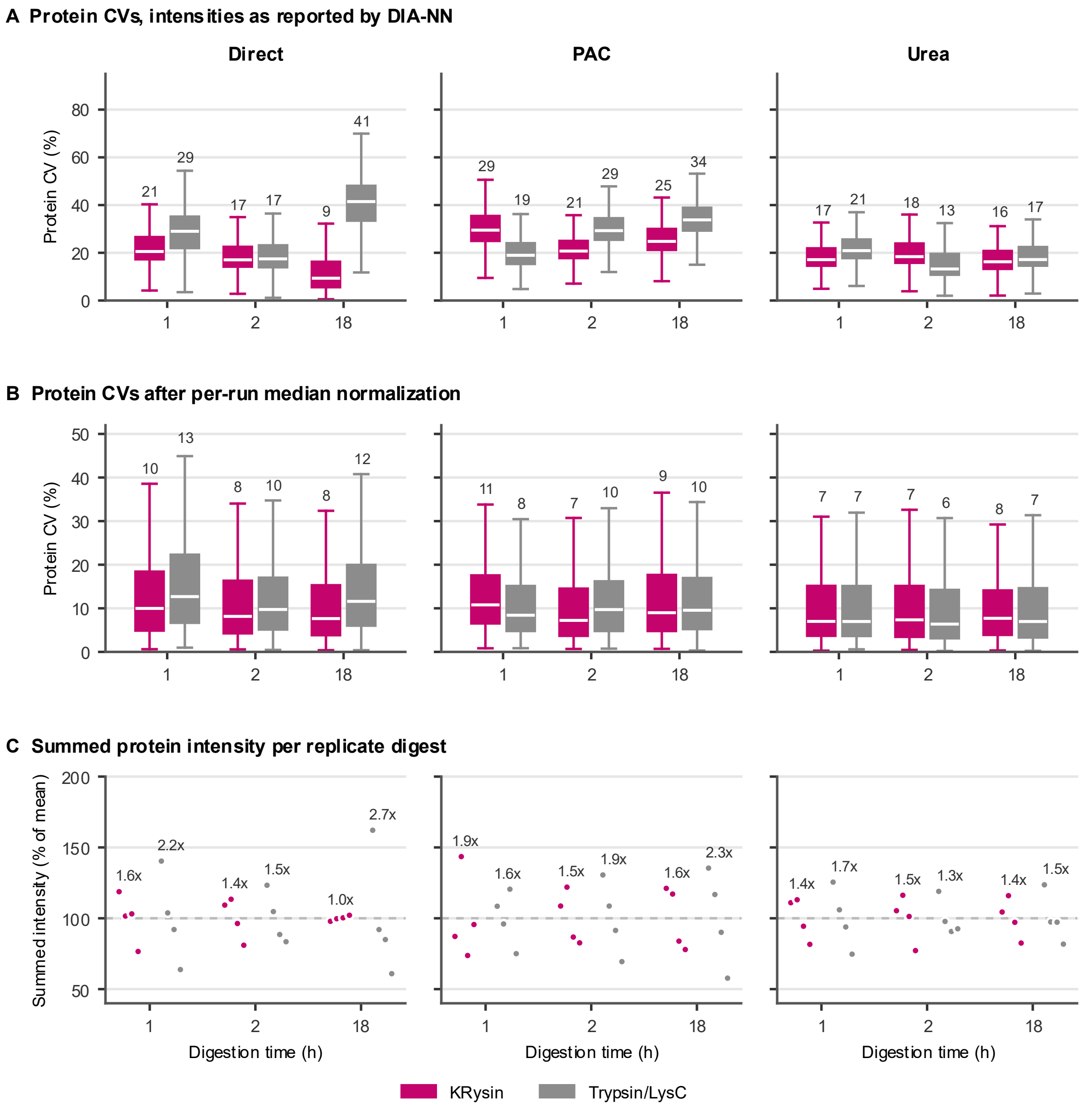

(A) Protein coefficients of variation (CVs) across four replicate digests for the direct, PAC and urea sample preparation workflows after 1, 2 and 18 hours of digestion, calculated from unnormalized DIA-NN protein group intensities for protein groups quantified in at least three of four replicates. (B) Protein CVs for the same data after per-run median normalization. (C) Summed protein group intensity of each replicate digest, shown as a percentage of the mean for that condition. Numbers above each condition give the ratio between the highest and lowest replicate. In A and B, boxes show the median and interquartile range, whiskers extend to 1.5 times the interquartile range, and median CVs are labelled above each box. KRysin is shown in magenta and trypsin/LysC in gray.

**Supplementary Figure S13. Protein quantification reproducibility as a function of protein abundance for KRysin and Trypsin/LysC digestion of human platelet-poor plasma.**

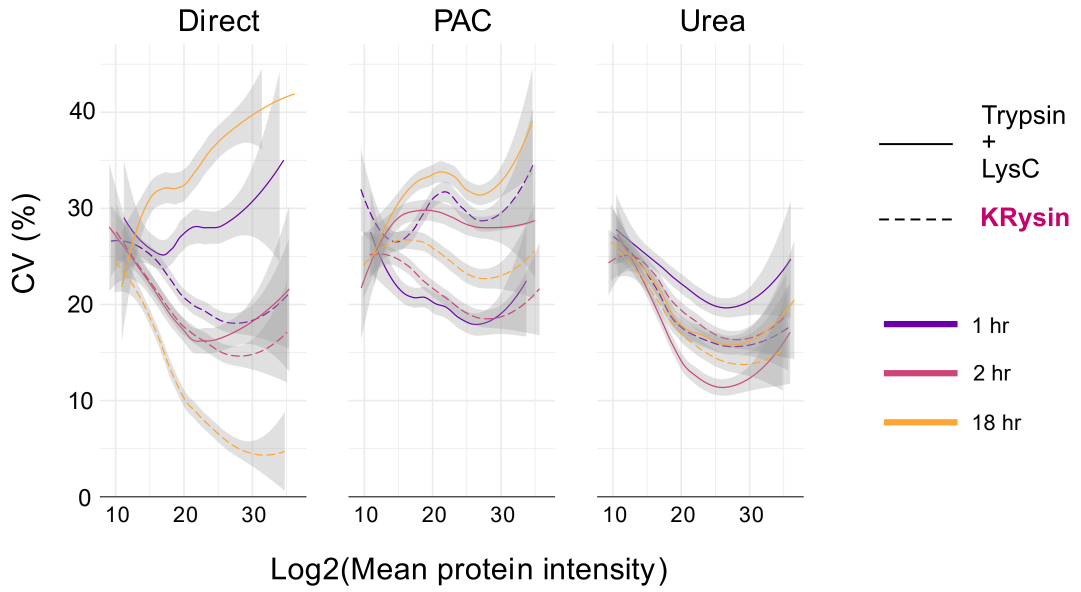

The coefficient of variation (CV, %) of protein quantification across replicate digests is plotted against log2 mean protein intensity, measured by narrow-window DIA. Human platelet-poor plasma was digested with KRysin (dashed lines) or Trypsin/LysC (solid lines) for 1 hour (purple), 2 hours (pink), or 18 hours (orange) using the Direct, PAC, or Urea workflow (panels). Lines show smoothed trends of the protein CVs, and shaded areas show the confidence interval of the fit. At mid to high abundance, KRysin showed comparable or lower CVs than Trypsin/LysC across the three sample preparation workflows.

#### Supplementary Figure S14. Digestion of human platelet-poor plasma with KRysin or Trypsin/LysC analyzed by data-dependent acquisition on an Orbitrap Exploris 480.

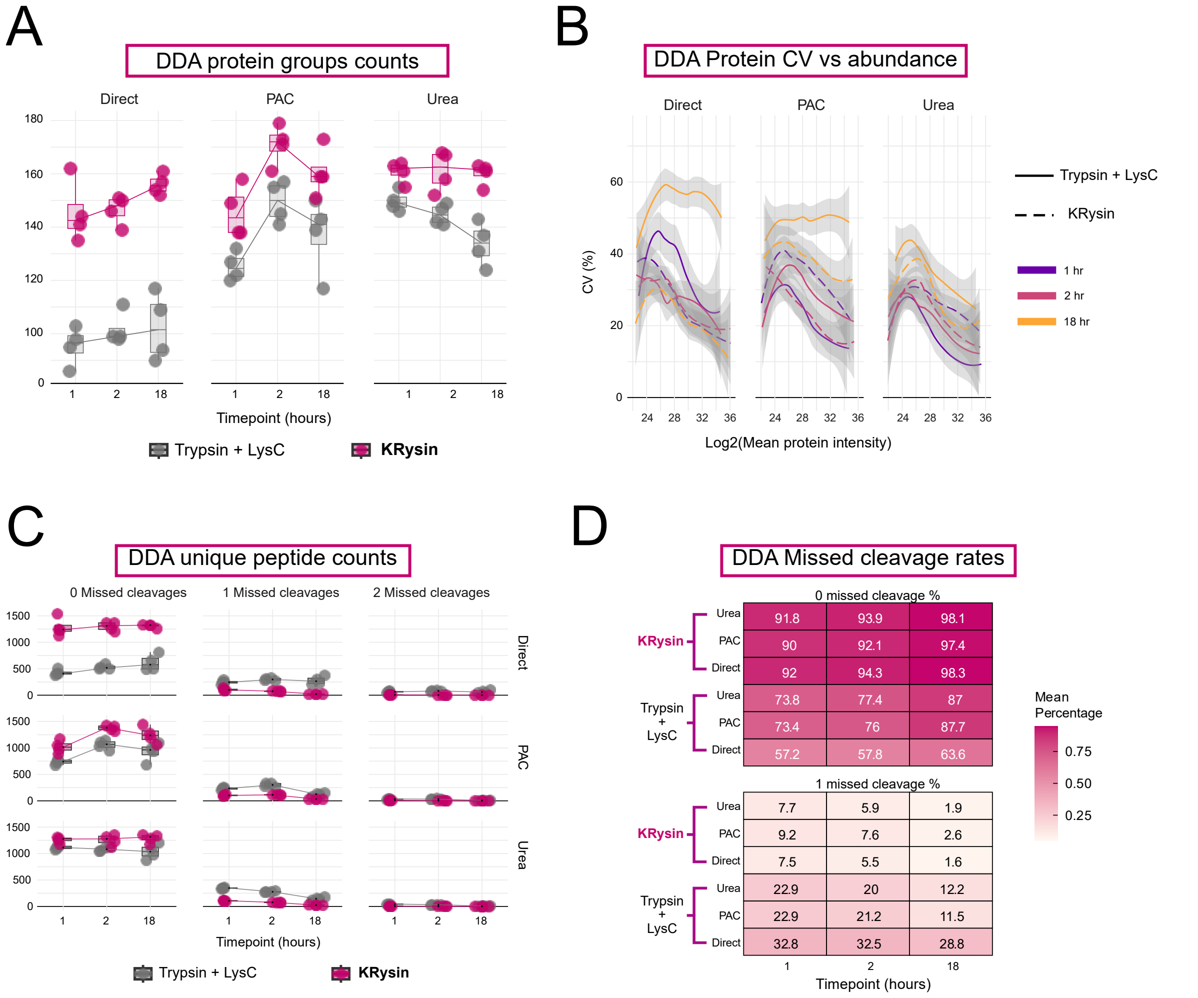

The same plasma digests tested with different sample preparation methods analyzed by narrow-window DIA (Figure 4E) were analyzed by LC-MS/MS using a data-dependent acquisition (DDA) method on an Orbitrap Exploris 480 mass spectrometer. Human platelet-poor plasma was digested with KRysin or Trypsin/LysC for 1, 2, or 18 hours using the Direct, PAC, or Urea workflow. (A) Number of protein groups identified. Points represent individual replicate digests, boxes indicate the interquartile range, and lines connect the medians across timepoints. KRysin yielded more protein groups than Trypsin/LysC in all three workflows. (B) Number of unique peptides identified with 0, 1, or 2 missed cleavages. KRysin yielded more peptides without missed cleavages in the Direct and PAC workflows, comparable numbers in the Urea workflow, and fewer peptides with one missed cleavage in all workflows. (C) Mean percentage of unique peptides with 0 (top) or 1 (bottom) missed cleavage. KRysin is shown as dashed lines and Trypsin/LysC as solid lines, colored by digestion time.
